# Label-free observation of individual solution phase molecules

**DOI:** 10.1101/2023.03.24.534170

**Authors:** Lisa-Maria Needham, Carlos Saavedra, Julia K. Rasch, Daniel Sole-Barber, Beau S. Schweitzer, Alex J. Fairhall, Cecilia H. Vollbrecht, Brandon Mehlenbacher, Zhao Zhang, Lukas Tenbrake, Hannes Pfeifer, Edwin R. Chapman, Randall H. Goldsmith

## Abstract

The vast majority of chemistry and biology occurs in solution, and new label-free analytical techniques that can help resolve solution-phase complexity at the single-molecule level can provide new microscopic perspectives of unprecedented detail. Here, we use the increased light-molecule interactions in high-finesse fiber Fabry-Pérot microcavities to detect individual biomolecules as small as 1.2 kDa with signal-to-noise ratios >100, even as the molecules are freely diffusing in solution. Our method delivers 2D intensity and temporal profiles, enabling the distinction of sub-populations in mixed samples. Strikingly, we observe a linear relationship between passage time and molecular radius, unlocking the potential to gather crucial information about diffusion and solution-phase conformation. Furthermore, mixtures of biomolecule isomers of the same molecular weight can also be resolved. Detection is based on a novel molecular velocity filtering and dynamic thermal priming mechanism leveraging both photo-thermal bistability and Pound-Drever-Hall cavity locking. This technology holds broad potential for applications in life and chemical sciences and represents a major advancement in label-free *in vitro* single-molecule techniques.

## Introduction

Tools to measure the properties of individual molecules (*1, 2*), including in heterogenous solutions (*3–9*), have become cornerstones of modern molecular and biomolecular research. Nearly all single-molecule approaches use extrinsic labels, and while these labels provide important contrast and specificity (*10*), label-free approaches that avoid arduous dye labeling procedures which may perturb the native functionality of biomolecules (*11–13*) are an increasingly desirable alternative. Most single-molecule approaches, including all current label-free methods, also rely on surfaces for immobilization, which is a costly compromise, as the measurement may bias detection towards sub-populations in mixed samples, disrupt native molecular interactions, alter dynamics, and generally precludes quantifying valuable solution-phase properties such as the diffusion constant (*9, 14, 15*). Here, we report a label-free single-molecule technique that enables detection of small solution-phase biomolecules (down to 1.2 kDa) with unprecedented signal-to-noise ratio (SNR) and allows resolution of their diffusion behavior.

Many label-free single-molecule experiments take the form of molecular detection, whereby the presence of a single copy of a specific molecule is perceived, typically through the presence of a surface-bound, selective, tight binder like an antibody. Other approaches take the form of molecular property assays, and extract information about the molecule, such as location, mass, or spectroscopic profile. Property assays are typically incapable of unambiguously identifying the molecule but can be applied generally, whereas molecular detectors can provide selective identification, but only for a small subset of chosen molecules.

The gamut of label-free single-molecule technologies has grown substantially, particularly across two modalities: interference-based and optical microcavity-enhanced techniques. Interferometric measurements, which generally rely on interference between elastically scattered light and a local oscillator, can operate as molecular detectors (*16, 17*) or property assays capable of determining position and mass (*18–22*). Dielectric optical microcavity platforms provide enhanced light-matter interactions due to high quality factor (Q) and low mode volume (V) (*23–25*). Microcavities have most commonly been applied as molecular detectors, where plasmonic enhancement (*26–28*), optomechanical coupling (*29*), or computational noise suppression (*30*) have enabled single-molecule detection via the reactive sensing mechanism (*31*) in which the interaction between a cavity mode and a molecule introduces a shift in the resonance frequency. Microcavities can also be used as single-particle property assays providing details on size (*32, 33*) or spectral information on electronic (*34*), plasmonic (*35*), or vibrational properties (*36*), and dynamics (*37*).

However, these approaches require target molecules to be surface-immobilized to allow signal integration and background subtraction, be bound by a surface-supported selective binder, or require interaction with a surface to couple to evanescent modes. The requirement for molecule-surface contacts can introduce perturbations to the native behavior (*9, 14*) while also obscuring solution-phase properties and dynamics. Open-access Fabry-Pérot microcavities can mitigate these concerns by operating in solution (*38*) but have not reached the single-molecule label-free regime. Recently, high-finesse fiber-based Fabry-Pérot microcavities (FFPCs) (*39*) were applied as effective sensors of single diffusing solution-phase silica nanoparticles (*40*). By extracting the frequency shift of optical modes, the formation of higher-order spatial modes, and the change in transmission intensity, the nanoparticle position was tracked, and the subsequent diffusion information was calculated. Here, we take advantage of the open-access geometry of FFPCs to achieve sensing of single, freely diffusing small biomolecules. Our platform utilizes passive mechanical stabilization of FFPCs (*41*), a new dynamic thermal priming mechanism, and active resonance frequency-stabilization as a novel form of molecular velocity filtering, to achieve detection of a 10 amino-acid, 1.2 kDa solution-phase single-protein with SNR of up to 123.

This observation is achieved in the absence of external surface-based signal multipliers like plasmonic enhancement and is the highest SNR reported for label-free single-molecule sensing by a substantial margin. Most importantly, the method operates without interaction with surfaces, allowing interrogation of unperturbed label-free solution-phase molecules, and evaluation of molecular diffusion profiles, a carrier of key information on biomolecule conformation and binding (*42*).

## Results

The FFPC was assembled from two single-mode optical fibers with concave laser-ablated end facets (Fig S1) which were subsequently coated with high-reflectivity dielectric layers (supplementary information) (*39*). The fiber mirrors were aligned and affixed laterally within a cut fused silica ferrule (Fig 1A, B) to increase the passive mechanical stability of the resonator (*41*). The optical modes were probed with static-frequency lasers over 660-760 nm, with laser output injected into the input fiber. Reflection and transmission channels were independently monitored on a pair of photodiodes (Fig 1A). The mirror separation was approximately 20 μm (Fig 1B), leading to Q-factors of ∼2×10^6^ and mode volumes on the order of 80 μm^3^ (*39*). The cavity finesses ranged from 27,000-101,000, across multiple cavities (supplementary information), in ambient conditions, reducing to 17,000-37,450 in water (Fig 1C). Continuous probing of a single resonant mode was achieved via phase-sensitive Pound-Drever-Hall (PDH) frequency locking (*43, 44*), in which the cavity length was actively stabilized to a single frequency of the pump laser (Fig 1A).

**Figure 1.**
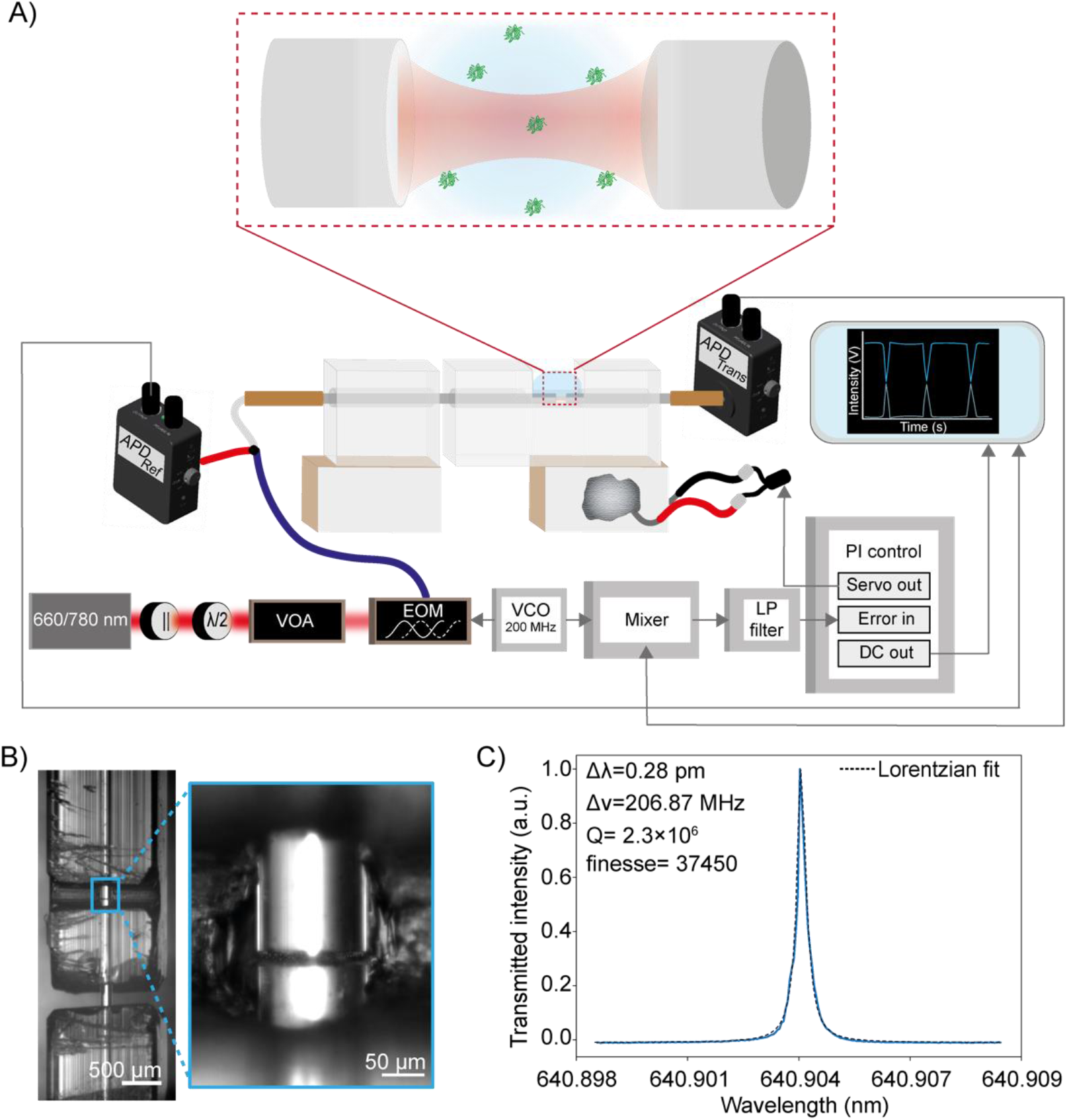
**A)** Simplified schematic of the FFPC-based single-molecule sensing instrumentation. Laser light (660-760 nm) of <1 MHz spectral width was transmitted through a linear polarizer (||) and half-wave plate (λ/2), selectively attenuated with a variable optical attenuator (VOA), and phase-modulated through a lithium niobate electro-optic modulator (EOM) driven by a 200 MHz voltage-controlled oscillator (VCO). Light was then coupled into the cavity via a fiber splitter, to enable collection of reflected light, and into an input optical fiber with transmitted intensity detected on a photodiode. PDH cavity-length stabilization, in order to maintain the cavity on resonance with the laser, was achieved using the frequency sidebands generated by the EOM driven by the VCO at 200 MHz. The error signal was generated by applying a low-pass filter to the mixed VCO reference and photodiode signals. This signal was then fed into the proportional-integral (PI) controller, which drives the ceramic piezo actuators to stabilize the cavity length to maintain resonance. Protein diffusion events were monitored in two channels on separate photodiodes, reflection and transmission. **B)** Brightfield images of the FFPC optical fibers within the quartz ferrule. The fibers were affixed within the ferrule, forming a cavity 19.3 μm in length. **C)** Wavelength scan used to determine the spectral linewidth of the cavity modes to be probed. The cavity finesse was 37450 in water.

We demonstrate the ability to detect single label-free proteins and small peptides by introducing samples of varying mass and radius into the FFPC. These included tetrameric streptavidin (66 kDa, 2.80 nm) (*45*), carbonic anhydrase (30 kDa, 2.10 nm) (*46*), aprotinin (6.5 kDa, 1.45 nm) (*47*) and c-Myc peptide, known more commonly as Myc-tag (1.2 kDa, 0.75 nm) (*48*). Protein samples were prepared at pM concentrations such that the mean occupancy of the optical mode volume was much less than one molecule. The input power into the cavity was ∼5 μW resulting in a circulating power of 5.5 mW.

Intensity traces show high amplitude, correlated signals in both transmission and reflection detection channels from transient interactions between single diffusing protein molecules and the locked cavity mode (Fig 2A), manifesting as a negative peak in transmission and a positive peak in reflection (mechanism discussed below). Confirmation that perturbation of the locked cavity originated from biomolecule diffusion and not ambient noise was achieved with water background measurements taken before the introduction of the protein and after removal, during which no signal was observed (Fig S2) and showing that detected events increased linearly with protein concentration (Fig S3). Time traces were recorded in 30-second intervals (Fig S4) with a temporal resolution of 20 μs, with the temporal scale of the single protein diffusion events on the order of 1-2 ms (Fig 2B, Fig S5). The extraordinarily high SNRs of up to 123 for Myc-tag (Fig S6) facilitated high temporal resolution with diffusion events able to be observed with at least 50 kHz sampling rate.

**Figure 2.**
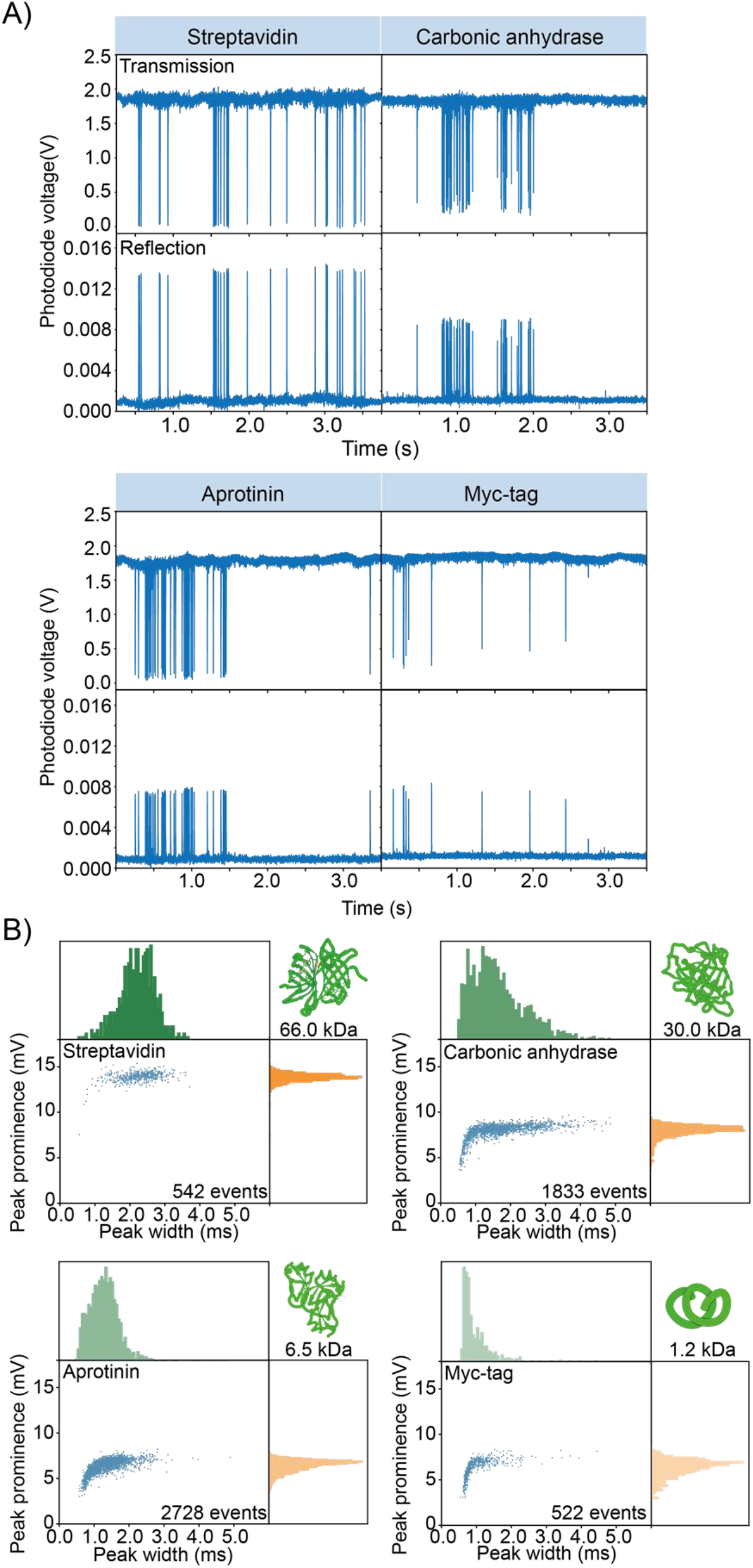
**A)** Perturbations of the locked resonant cavity mode originating from single-protein diffusion events. The locked signal was monitored in both transmission and reflection, and the events manifested as a transient reduction of the transmitted signal intensity and an increase in the reflected intensity. **B)** 2D plots and accompanied histograms of the extracted prominences and temporal widths of the reflected signals. The corresponding transmission data can be found in Fig S7.

Without relying on plasmonic enhancement mechanisms (*26–28, 49*), surface proximity (*17, 26–30, 49*), or the consequent conformational or chemical change of a surface-supported docking molecule (*30, 49*), we demonstrate SNRs of up to 42-fold higher than existing label-free biomolecule sensing techniques for molecules of comparable molecular weight (*17, 26*–*30*) as well as achieving a mass limit of detection ∼25-fold smaller than that of mass photometry (*20*).

Each transit event comprises both temporal and intensity data. Plotting the distribution of temporal and intensity parameters provides a 2D distribution signal profile containing unique information on the molecular mass and diffusion (Fig 2B). Each protein molecule exhibited a distribution of temporal widths, identified from the full width at half maximum (FWHM) of events that rise significantly above the noise, and prominences, which increased with increasing protein molecular weight (Fig 2B). A diversity of widths is expected due to the stochasticity of Brownian motion. The prominence of the peaks differed between transmission and reflection detection channels (Fig 2B, Fig S7); this behavior arises from the dispersion that is unique to FFPC cavities (Fig S8) (*50*). The mean temporal widths of the events were unchanged at proportional gain values > -50 dB in the proportional-integral (PI) control of the PDH system (Fig S9), where a higher proportional gain value constitutes a higher locking bandwidth (LBW). Consequently, experiments were conducted above this threshold at a LBW of ∼5 kHz (Fig S10). Taken together, these data confirm that these high amplitude signals originate from the perturbation of the cavity mode volume by single diffusing proteins.

To demonstrate the ability of this technique to move beyond simple detection, we explored the potential for property assay using correlation analysis to extract temporal information from the data. Correlation spectroscopy is a ubiquitous tool across temporally sensitive biophysical methods, such as fluorescence correlation spectroscopy (FCS) and dynamic light scattering, aiming to extract ensemble diffusional and, therefore, size and mass information of molecules (*42, 51–55*). The autocorrelation expectedly shows dynamics on longer timescales for proteins of increasing mass (Fig 3A). Importantly, the autocorrelation times were consistently linear in proportion to the radius of the protein (Fig S11). This result confirms the versatility of this new single-molecule technique as a molecular property assay, demonstrating the potential to extract meaningful molecular information, including size and diffusional properties. Label-free methods of assessing molecular dynamics can offer substantial impact in biophysical applications.

**Figure 3.**
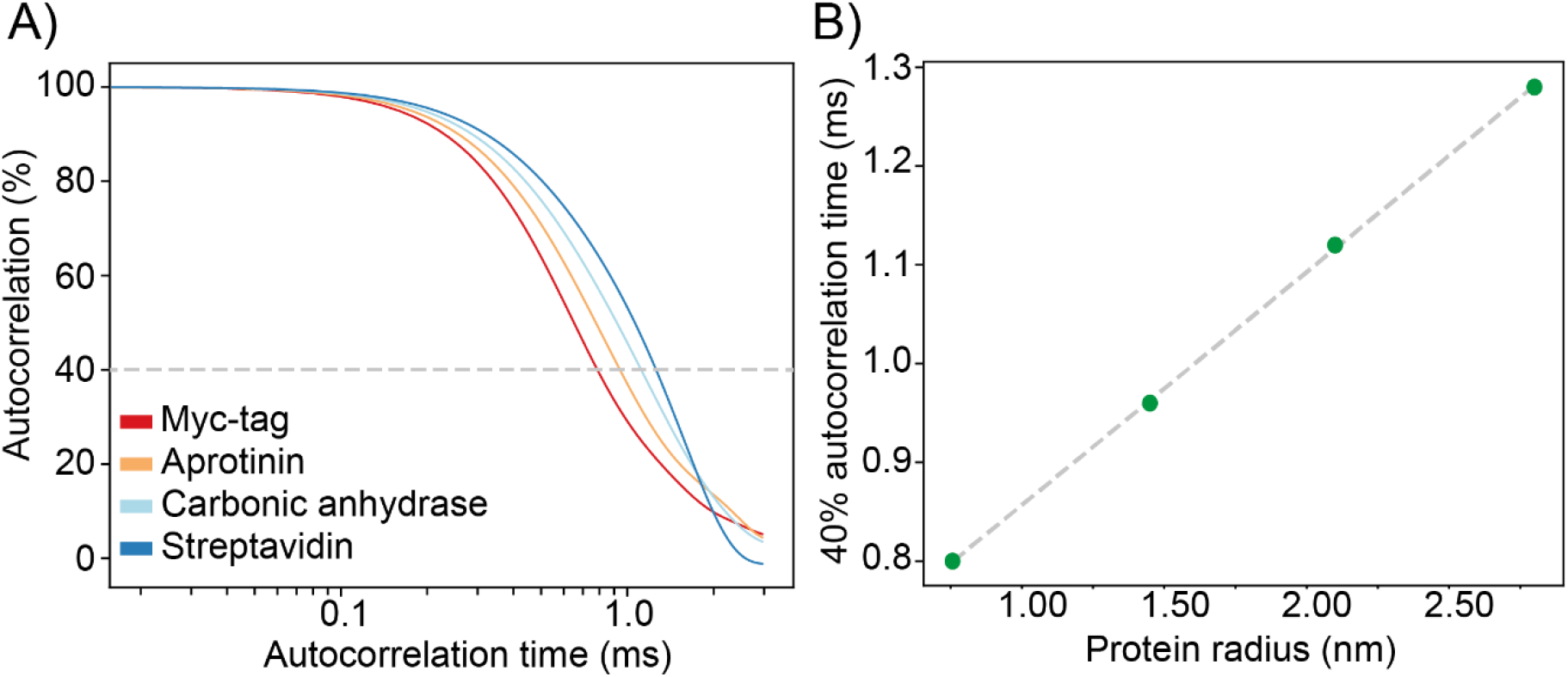
**A)** Ensemble autocorrelation of several hundred single-protein diffusion events. **B)** Relationship between autocorrelation time at an autocorrelation threshold of 40% (see Fig. S11 for other thresholds) and the protein radius, showing a clear linear correlation.

The extraordinarily high SNR of the single protein events, even down to the 10 amino-acid peptide Myc-tag, highlights the potential to extend the dynamic range of applications to both smaller molecules and higher acquisition rates. To demonstrate this, we measured transient events of Myc-tag diffusion with 2 μs time resolution (500 kHz acquisition rate, Fig S12). This high collection frequency facilitated the measurement of events as narrow as 26 μs. With a 500 kHz collection frequency, even these high-speed events can be sampled far beyond the minimum Nyquist requirement, highlighting the potential to study kHz processes such as enzyme kinetics and conformational changes (*56*) without sacrificing SNR. Only the photothermal bandwidth will ultimately limit the temporal resolution (see below).

Having demonstrated single-molecule measurements of solution-phase, label-free proteins, we next demonstrated the ability of our system to resolve populations of simple bimolecular mixtures. Resolution of mixtures is vital for identifying diagnostic biomarkers, understanding disease pathogenesis, and for elucidating biomolecule-biomolecule and biomolecule-drug interactions. Techniques such as FCS, invaluable for inferring conformation, and fluorescence polarization anisotropy, invaluable for ascertaining drug binding (*57*), are limited by the requirement for fluorescent labels. Ensemble label-free techniques such as dynamic light-scattering can provide diffusive information of biomolecules, but analysis is restricted by the high dependence of scattering on the molecular radius (r^6^), obscuring small particles among larger ones (*58*), necessitating monodisperse samples for quantitation. Mass photometry overcomes this obstacle via spatial discrimination but is constrained by limit of detection and use of surfaces (*20*). Our approach results in 2D profiles that can act as molecular signatures (Fig 2B) containing information about mass and diffusivity.

First, we investigated a mixture of aprotinin and Myc-tag, which have a 5.3 kDa mass and 0.7 nm radius difference (Fig 4A). The 2D profile of the mixture is qualitatively similar to the component distributions. Though these two populations would be difficult to resolve considering only peak prominence, two distinct populations, a fast-moving population with a mean event FWHM of 0.49 ± 0.15 ms and a broader, slow-moving population with a mean event FWHM of 1.68 ± 1.37 ms, are clearly evident.

**Figure 4.**
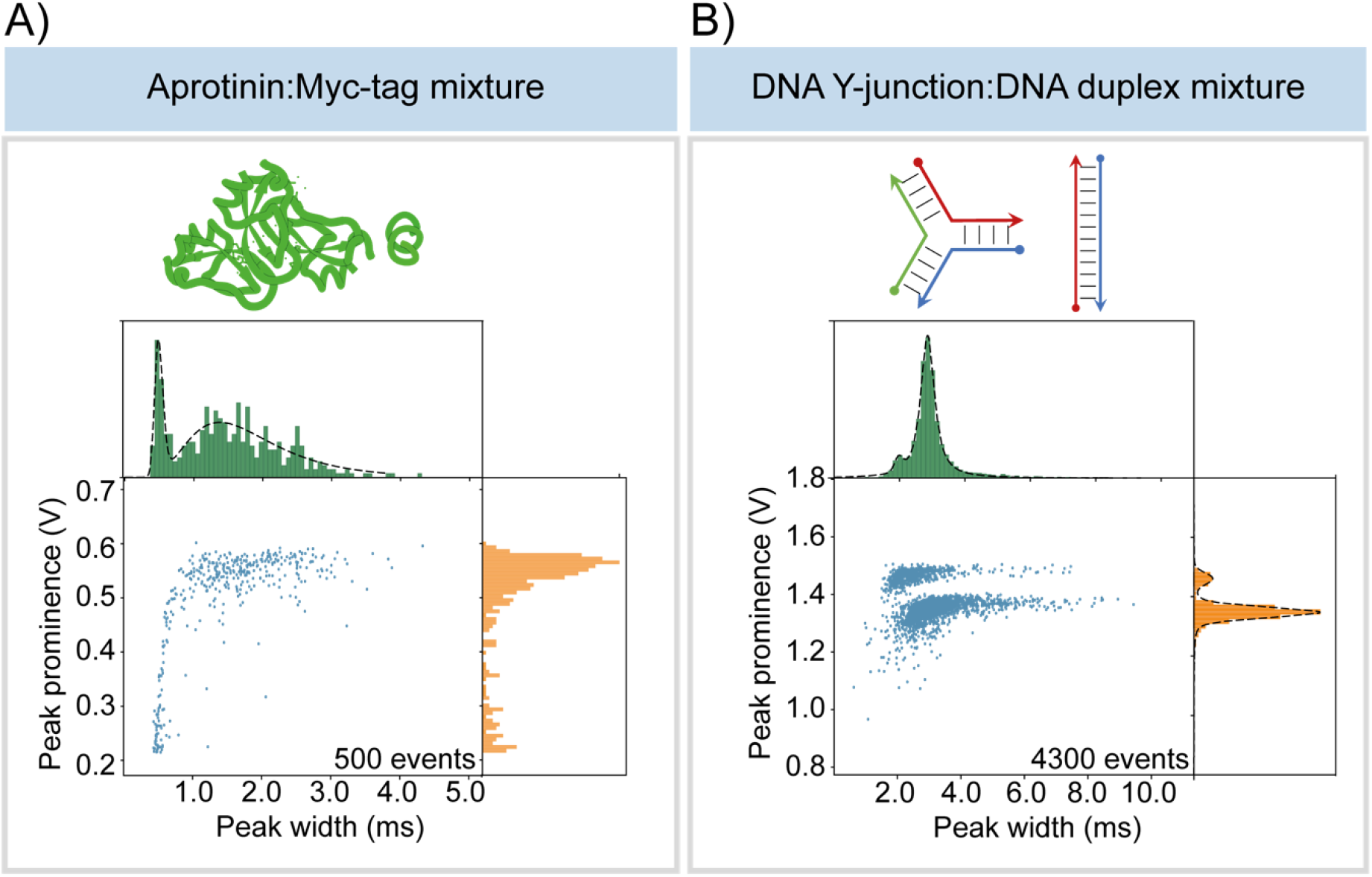
2D plots of peak prominence versus temporal width and subsequent independent histograms for **A)** a mixed protein sample of aprotinin (6.5 kDa, 1.45 nm) and Myc-tag (1.2 kDa, 0.75 nm) and **B)** a mixed DNA structure sample of a duplex (16.6 kDa, 9 nm) and Y-junction (16.6 kDa, 5 nm), with multiple populations clearly resolved.

Moving beyond protein samples, we explored the resolution of bimolecular mixtures of DNA isomers of identical mass (16.6 kDa) and composition, but differing sequence (Fig 4B): a DNA duplex (9 nm) and a Y-junction structure (5 nm). Here, the two populations are clearly resolved in both dimensions of the 2D profile. The event prominence was distinctly separated into two populations, a low-intensity population with a mean prominence of 1.35 ± 0.02 V and a lower abundance high-intensity population with a prominence of 1.46 ± 0.03 V. Two partially resolvable populations of similar relative magnitudes to those observed in peak prominence were visible in the temporal domain. Interestingly, the more rapidly diffusing component of the mixture produced a larger magnitude perturbation to the cavity mode. As discussed below, the response of the FFPC to the molecular perturbation is influenced by molecular properties as well as multiple dynamic cavity properties. The ability to cleanly reveal the presence of two molecules of identical small mass but differing conformation and diffusion behavior shows that this approach provides complementary information not discernable from mass photometry.

## Discussion

The detection of a single, freely moving, un-labeled small biomolecule with high SNR requires a plausible mechanism whereby the small perturbation to the optical system can be discerned. Our proposed mechanism begins with a refractive index change as the biomolecule displaces water molecules of lower index in the microcavity (often referred to as the “reactive mechanism”) (*31*). Resonance shifts of 1-49 kHz due to the altered optical path length are estimated from the protein molecular weights (Fig S13). We note that these shifts are ∼20× greater at equivalent weights than estimates in whispering gallery mode resonators (*30*) due to smaller mode volume and better spatial overlap between molecule and optical mode in FFPCs (supplementary materials). The ability to resolve resonance shifts that are small compared to the cavity linewidth (∼200 MHz) with such high SNR is based on a combination of high passive stability, active low-frequency stabilization, creation of a velocity discrimination window for molecular motion, and the use of dynamic photothermal distortion of the resonance line shape. In water the photothermal effects occur due to absorption of some of the cavity circulating power which alters the refractive index of the medium via the thermo-optic coefficient.

The combination of mounting the FFPC in a glass ferrule and PDH locking provides remarkable stability to the optical system, suppressing mechanical and laser frequency noise to detector-limited levels (Fig 5A) (*41*). Furthermore, the mechanical stability of the cavity is extremely high, well-below the detector noise floor (Fig S14). Importantly, the PDH LBW only suppresses fluctuations (including molecular fluctuations) at temporal frequencies below 5 kHz (Fig 5A). This is a critical function of the PDH loop, as low-frequency mechanical fluctuations can introduce substantial resonance frequency shifts. This loop would also suppress perturbations produced by larger, slow-moving molecules or particles, as the majority of their displacement would occur within the PDH LBW (Fig 5A). Importantly, small molecules undergoing Brownian motion, albeit with smaller overall resonance shifts due to their reduced size, have a larger fraction of their mean squared displacement power spectral density (MSDPSD) (*59*) outside the PDH suppression window (Fig 5A, Fig S15).

**Figure 5.**
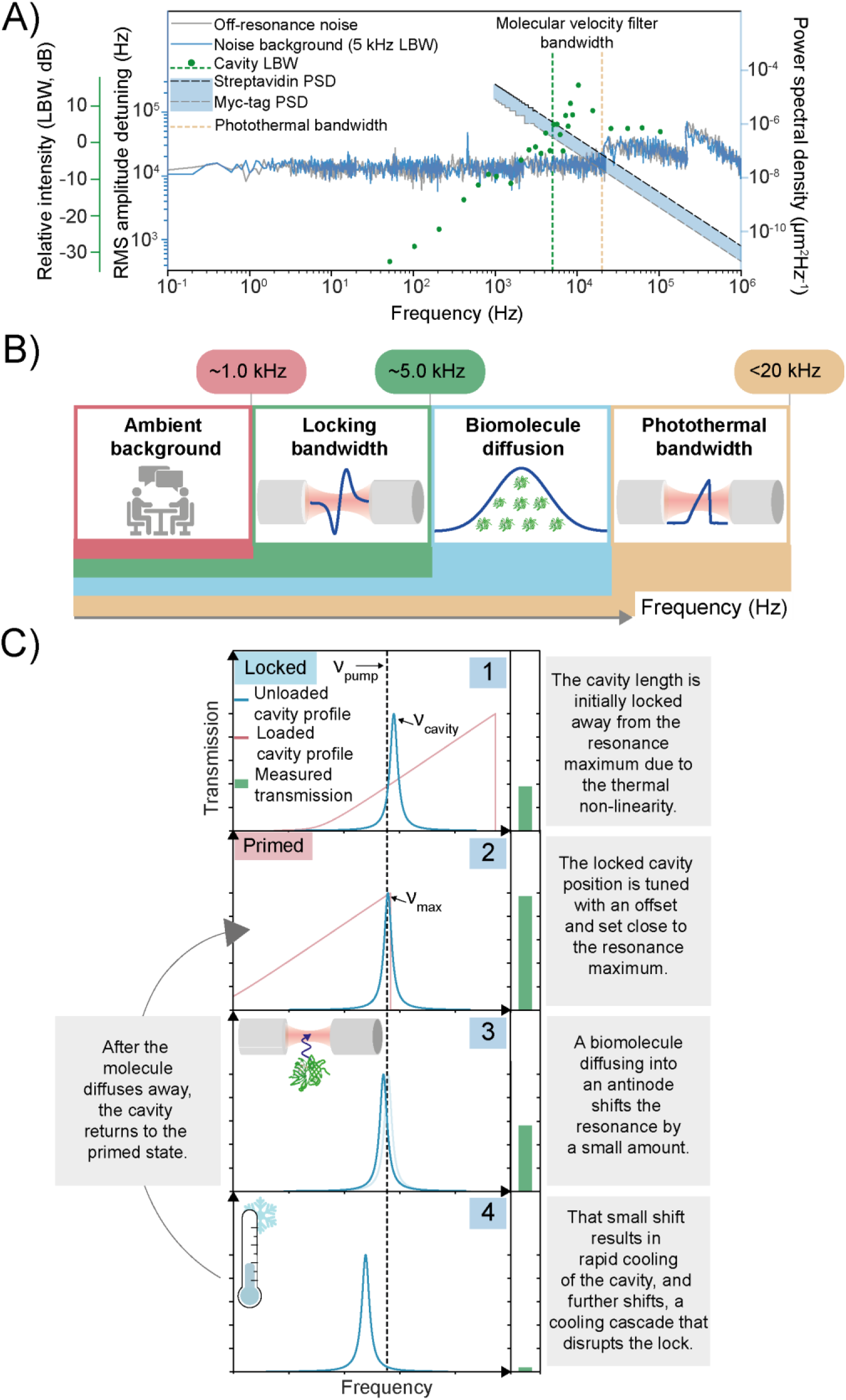
**A)** Plot showing frequency noise spectral density in water, LBW characterization, and mean-square-displacement power spectral density (MSDPSD) of proteins, streptavidin, aprotinin, carbonic anhydrase, and Myc-tag. The noise spectral density of the locked cavity in water rapidly converges to the detector-limited noise (off-resonance noise), highlighting the high passive stability. The locking bandwidth of 5 kHz, defined by the 0 dB feedback gain crossing, governs the lower frequency limit of the velocity filter. The upper limit of the velocity filter is defined by the photothermal bandwidth (150 kHz). The molecular MSDPSD can be integrated within this filter bandwidth to determine the root-mean-square MSD **B)** Cartoon illustrating the key processes and their frequency bandwidths. Noise below 5 kHz is suppressed by the PDH. The upper limit of the LBW and the lower limit of the photothermal bandwidth define the molecular diffusion velocity observation window. **C**) Schematic describing the mechanism of dynamic thermal priming (see text for details).

Integration of the MSDPSD for the smallest protein (Myc-tag, 0.75 nm) between the end of the locking bandwidth and the photothermal bandwidth (discussed below) yields a root-mean-squared (RMS) displacement of 93 nm (supplementary materials). This displacement is comparable to the ∼250 nm distance between the node and antinode of the cavity standing wave (Fig S16), suggesting that the near full resonance shift can be experienced by the microcavity outside the PDH LBW due to diffusing molecules. When the molecule diffuses back into the node, the perturbation ceases, leading the system to exhibit a dependence on the molecular diffusion constant (Fig 3). Detection by shifting from node to antinode is distinct from the operational mechanisms in evanescent detection modalities (*17, 27, 28, 30, 49*). Critically, a solution-phase label-free apparatus allows this novel employment of PDH as a high-pass filter to reduce mechanical noise while passing signals from fast-moving molecules (Fig 5B), a key difference compared to previous schemes.

The second element of our proposed mechanism relies on a photothermally induced distortion of the resonance line shape (Fig 5C) and a dynamic photothermal priming mechanism, which amplifies small resonance shifts. FFPCs spatially confine relatively intense optical fields, inducing on resonance temperature changes inside the mode volume media and consequent thermo-optic resonance shifts (*60*). The presence of this thermal nonlinearity clearly manifests as a broadened asymmetric cavity line shape upon active scanning of the cavity length or wavelength (Fig S17) (*61*). This photothermal nonlinearity arises from the mirror coatings and the aqueous medium and requires no light absorption by the molecule itself (*31, 62*). The photothermal bandwidth was determined experimentally to be 21 kHz (Fig S18A), defining the upper limit of our molecular observation window (Fig 5B). In a non-PDH stabilized cavity, these photothermal nonlinearities result in multiple distinct stable equilibria (*60*). However, with PDH stabilization, this nonlinearity can be used for additional signal amplification. Increasing the cavity transmission by introducing an offset from the original, arbitrary locked position (Fig 5C, panel 1) results in the pump laser sitting at a frequency just lower than the cavity maximum (Fig 5C, panel 2). In this primed state, even the resonance shift of a diffusing molecule can shift the cavity resonance to an unstable regime where the pump laser sits at a higher frequency than the microcavity resonance (Fig 5C, panel 3). Here, the shift triggers a dynamic process by which the cavity cools faster than the LBW (Fig S18), resulting in further resonance shift and more cooling, ultimately causing a substantial transmission decrease (Fig 5C, panel 4). Other molecule-induced mechanisms, such as scattering, may also contribute to cavity cooling. After the molecule has diffused out of the antinode, the cavity begins to warm, and eventually, the PDH recovers the initial locked position at a rate defined by the LBW (Fig 5C, panel 2). In the case of smaller perturbations (as with Myc-tag), the cavity cools less, leading to the distribution of peak prominences. Evidence for this mechanism can be found in controlled voltage pulses added to the output servo of the PDH, providing internal perturbations (Fig S19A) that qualitatively mimic molecular passages (Fig S19B).

In summary, our proposed mechanism features molecules diffusing into the microcavity, where their fast motion exceeds the PDH locking bandwidth. A hypersensitive photothermally primed cavity, experiencing these fast molecular perturbations, rapidly cools, leading to enhanced shift and massive signal. Both peak prominence and temporal width are expected to be influenced by system parameters, including PDH LBW. However, while peak prominence is a complex function of biomolecule molecular weight (and thus refractive index) and diffusive parameters, the temporal width is expected to be dominated more purely by diffusive parameters, leading to clear linear dependence (Fig 3). Future work will allow more quantitative information to be derived from peak prominence.

## Conclusion

In the absence of surfaces, extrinsic labels, and plasmonic enhancers, this work has demonstrated exceptional sensitivity in observing single, diffusing biomolecules, achieving SNRs of >100 for a sub 1 nm peptide. Our approach leverages the open-access geometry of micro-scale FFPCs to facilitate unimpeded biomolecule diffusion as well as maximize the overlap between the biomolecules and the optical field. Our enhanced sensitivity relative to other label-free techniques originates in molecular velocity filtering and photothermal priming, where two experimental challenges, fast molecular motion, and thermal non-linearity, are transformed into advantages. Much like the fingerprint region of an infrared spectrum, the resulting rich 2D intensity/temporal data can be used to distinguish unique, identifying molecular signatures and has the potential to provide quantitative mass and diffusional information without surface perturbation.

Mass photometry, a new method that can provide quantitative mass information of unlabeled biomolecules in a spatially resolved manner (*20*), has been commercialized and widely adopted, showcasing the tremendous possibilities of photonic single-molecule assays. Our approach sacrifices the spatial resolution of mass photometry. On the other hand, our solution-phase FFPC-based approach avoids surfaces while providing μs dynamics, a substantially higher sensitivity with ≤ 1 kDa detection limit, and 2D signal profiles that offer a path toward distinguishing molecules based on conformation, which influences diffusion properties, as well as just mass. In addition, we note that FFPCs offer convenient fiber optic integration and that molecules, after passing through the FFPC, could be readily interrogated via mass photometry, making the approaches truly complementary.

Further refinement, including simple experimental advances such as increased suppression of external noise sources, is expected to yield significant improvements, including the capability to detect biomolecules smaller than 1 kDa. Optimization of measurement parameters using a quantitative model will enable tuning of molecular profiles, for instance, a configuration of the bandwidth of the velocity filter to selectively collect information from different diffusional populations. Our FFPC approach has the potential to resolve rapid biomolecular conformation changes, elucidate self-assembly of small molecules in complex samples, and provide routes to rapid screening of protein-protein and protein-drug interactions. By being label-free and single-molecule, our method can mitigate some of the key experimental difficulties in FCS and dynamic light scattering, two widely applied biophysical techniques. This straightforward and readily scalable apparatus will bring numerous benefits to the fields of life and chemical sciences, such as trace analysis, separation science, mechanistic insights, and clinical diagnostics.

## Acknowledgements

This work was principally funded by the National Institute of Health (NIH, R01GM136981), with resonator construction supported by the Q-NEXT Quantum Center, a U.S. Department of Energy (DOE), Office of Science, National Quantum Information Science Research Center, under Award Number DE-FOA-0002253 and the National Science Foundation (NSF) Quantum Leap Challenge Institute for Hybrid Quantum Architectures and Networks, Award No. 2016136, and additional instrumentation development supported by the Center for Molecular Quantum Transduction and the Energy Frontier Research Center funded by DOE, Office of Science, BES under award DE-SC0021314, and by Schmidt Futures. L.-M. N. was partially funded by the European Union’s Horizon 2020 research and innovation programme under the Marie Skłodowska-Curie grant agreement No 886216. E.C. was funded by NIH (MH061876 and NS097362). H.P. was funded by the Deutsche Forschungsgemeinschaft (DFG, German Research Foundation) under Germany’s Excellence Strategy – Cluster of Excellence Matter and Light for Quantum Computing (ML4Q) EXC 2004/1 – 390534769. We thank Brendan Cullinane, Yulia Podorova, Blaise Thompson, and Tracy Drier.

