## Supplementary Information for "Label-free observation of individual solution phase molecules"

This PDF file includes:

Materials and Methods

Figures S1 to S22

Tables S1 to S4

Supplementary references

### Table of Contents

**Materials and Methods** ... 4

**Experimental setup** ... 4

Mirrored fiber production ... 4

Ferrule-assembly Preparation ... 5

Fiber cavity construction ... 6

Optical setup ... 7

Hardware control and data collection software ... 8

Sample preparation ... 9

**Experimental data collection** ... 10

Single biomolecule diffusion experiments ... 10

Locking bandwidth characterization ... 11

Noise background experiments ... 11

Voltage pulse experiments ... 12

Photothermal broadening and thermo-optic coefficient ... 13

Photothermal bandwidth measurement ... 14

**Data analysis** ... 15

Single-molecule diffusion and autocorrelation analyses ... 15

Locking bandwidth analysis ... 16

Signal-to-noise ratio determination ... 16

Photothermal bandwidth determination ... 17

**Simulations and calculations** ... 18

Calculated resonance shift ... 18

Molecular velocity distributions and molecular mean-square-displacement power-spectral-density ... 21

Simulated photothermal bandwidth determination ... 23

### Materials and Methods

#### Experimental setup

##### Mirrored fiber production

Copper-coated optical fiber with cladding diameter of 125 µm (IVG Fiber, Cu600) was first etched using nitric acid (70%). This etching procedure removed the copper coating, leaving a layer of carbon coating above the cladding. The carbon coating was removed with a small amount of diamond paste on cloth; however, this step was removed for later fiber production due to apparent contamination from the diamond paste. Fibers were flat-cleaved (<0.2-1°) using an automated cleaver (AFL Fujikura, CT-106) (*1*).

An 18 W $CO_2$ laser (Synrad, 48-1 KAM) generated the ablation laser beam, which was guided and modified with polarizing, phase retarding, and other associated reflecting and ZnSe focusing optics. An arbitrary waveform generator (Agilent, 33220A) controlled the beam characteristics, and a digital optical power meter head (Thorlabs, S314C) with an attached console (Thorlabs, PM100D) was used to measure the beam power. Optical fiber substrates were mounted atop a fiber clamp (Thorlabs, HFF003), which rested upon a 3-axis translation system (Thorlabs, MTS50-Z8; MTS50B-Z8; MTS50C-Z8), driven by a DC servo motor controller (Thorlabs, KDC101) secured to a long-range single-axis translator for inspection-ablation positioning (Thorlabs, LNR502(/M)) driven by a motor controller (Thorlabs, BSC201). A CCD camera (Thorlabs, CS165MU) coupled to a long working-distance microscope system (Navitar, 160-10) with a 20× magnification, 0.42 NA infinity corrected objective (Mitutoyo, 378-804-3) illuminated by a 635 nm LED source (Thorlabs, LEDD1B), was used to image the fiber position. Optimal ablation alignment was facilitated by illuminating the fiber core with an optical fault finder (VFLTOOL, HGB30). The laser was set to have a power of 0.28 W, and a shot time of 250 ms was controlled using a shutter in front of the fiber. These shot parameters generated fibers with radii of curvature (ROC) at the base of the ablation of 48 µm on average, and diameters, calculated by 2-σ of a Gaussian fit to the ablation profile, of 21 µm on average (Figure S 1).

The fibers were characterized with a ZYGO Interferometer. Two perpendicular slices of the surface profile were analyzed. For calculating the ROC, the center of the ablation was assumed to be the minimum. A polynomial fit was then performed, and stepwise derivatives were averaged to calculate the ROC. For the diameter, Gaussian fitting was performed, and the diameter was taken to be two standard deviations of the fit. The ellipticity of the ablation was calculated by comparing both the ROC and the diameter values for the perpendicular slices. The decentration of the ablation was also measured by coupling the fault finder through the core of the fiber, which was then compared to the center of the ablation. The decentration, ROC, diameter, and ellipticity were all used to determine the viability of a fiber.

The fiber substrates were coated commercially with wavelengths of maximum reflectivity at either 635 nm (LASEROPTIK GmbH, Germany) or 780 nm (LAYERTEC GmbH, Germany), in which alternating layers of $Ta_2O_5$ and $SiO_2$ were deposited using ion beam sputtering (IBS). These generated distributed Bragg reflector surfaces with transmission loss < 20 ppm, absorption loss < 10 ppm and scattering loss < 16 ppm.

#### Ferrule-assembly Preparation

The cavity bridge assembly was prepared using fused silica ferrules (VitroCom, 8×1.25×1.25 mm) with an inner bore of 131 μm, based on the procedure by Saavedra *et al* (*2*). The glass ferrule was cleaned using an air plasma cleaner (Harrick Plasma, PDC-001-HP) for 10 minutes. A thin layer of UV-curable glue (Dymax, 9037-F) was placed on two 150 V piezos (Thorlabs, PA4DG), which were aligned flat to the ferrule, ensuring that the direction of translation was aligned with the long axis of the ferrule. The glue was then cured using a UV light (Rolence Enterprise, Q6 UV). The assembly of ferrule and two piezos was then affixed to a glass block, which was approximately 20×7×3 mm in dimension. This assembly was then placed in an oven at 65-75 °C for 2-3 hours for additional curing of the glue. After this, wires were soldered onto the piezos. Two cuts were made in the ferrule in order to facilitate translation of the cavity length: one full-cut to separate the ferrule into two parts, maximizing the cavity length translation, and one half-cut to preserve fiber alignment along the inner bore. These cuts were made using diamond wire (Ø=125 μm) secured in a jeweler's saw. The full-cut was made off center and the half-cut was made close to the center of the ferrule. After the full-cut, a small piece of wire was placed in the bore to alert when the cut had reached the bore. During the cutting process the glass dust was removed using canned air.

The ferrule was cleaned post-cutting by first running the ferrule under deionized water for 5 minutes. Using a digital microscope to visualize, Millipore water was pipetted through the bore of the ferrule and a piece of optical fiber was run through the bore to remove any remaining glass dust. This process was repeated until no visible dust remained. The ferrule was then rinsed under Millipore water and few drops of methanol (≥ 99.9 %) were run through the ferrule, which was then dried thoroughly using a nitrogen gas line.

#### Fiber cavity construction

The previously described high reflectivity, mirror coated optical fibers were spliced to a connectorized patch cable (Thorlabs, P3-630Y-FC-2) using a fusion splicer (Fujikura, FSM-100P). The fibers were aligned using a 6-axis piezo actuated stage (Thorlabs, MAX602D). The input fiber was mounted to this stage using a tapered v-groove fiber holder (Thorlabs, HFV002). The output fiber was held by a v-groove fiber holder secured to an XYZ translation stage. To visually align the fibers, two cameras were aligned perpendicular to the fiber axis. The top-down view used a CMOS camera (Thorlabs, DCC1545M) connected to a zoom lens (Navitar, 1-50487). This imaging system was used for both alignment and to estimate the distance between the fibers. The second perpendicular axis was visualized using a digital microscope (Dino-Lite, AM4113ZT). Using both camera axes, the fibers were then coarsely aligned to one another visually using the 6-axis stage. The next step was to finely align the fibers by recording resonances. To do this, a ramp signal from a data acquisition board (DAQ, NI, BNC-2120) was applied to a piezo controller (Thorlabs,

MDT693B), which drove the piezo aligned with the fiber axis. Laser light (635-760 nm) was then injected into the input fiber and collected through the output fiber to an avalanche photodiode (APD, Thorlabs, APD430A). Resonances were measured in transmission and optimized by adjusting all parameters of the 6-axis stage. The cavity finesse was characterized by either using a wavelength tunable external cavity diode laser (Newport, TLB-6704) or via an electro-optic phase-modulator (EOM, EOSpace, PM-0S5-10-PFA-PFA-633, PM-0S5-01-PFA-PFA-765/781) by introducing sidebands at a known frequency (2.6 GHz) and the linewidth extracted via Lorentzian fitting.

The fibers were guided into the prepared ferrule using the top-down and perpendicular views and the cavity formed at the center of the half-cut. In order to secure the fibers inside the ferrule, 2 μL of low viscosity UV-curable glue (Masterbond, UV16) was deposited onto each fiber in turn at the left entrance of the non-slotted ferrule and the right entrance of the slotted ferrule. The glue was drawn up the fiber using capillary action and was UV-cured (Rolence Enterprise, Q6 UV) after ~2 mm of travel into the ferrule. The piezos on the bridge were then attached to a piezo driver (nPoint, D.200) and driven to assess the resonances under constant cavity translation.

#### Optical setup

Experiments were performed on a custom-built setup (Figure S 21). Either a single-frequency, continuous-wave diode laser (660 nm, Cobolt Flamenco, 90261, 300 mW) of < 1 MHz linewidth or a tunable Ti-Saph cavity laser of <100 kHz linewidth operating at a single frequency (760 or 780 nm, MSquared, SolsTiS) was passed through a linear polarizer and half wave-plate then fiber coupled into an electronic variable optical attenuator (Thorlabs, V600), then into an EOM (EOSpace, PM-0S5-10-PFA-PFA-633, PM-0S5-01-PFA-PFA-765/781) driven by a voltage-controlled oscillator (VCO, Mini-Circuits, ZX95-209-S+, 200 MHz). The light was then coupled into the input fiber of the cavity via a fiber-based beam splitter (Thorlabs, TW670R3A1), the power injected into the cavity was between 5-35 μW. The circulating power ($P_{circ}$) was calculated using Equation 1:

$$P_{circ} = P_{in}T\left(\frac{F^2}{\pi^2}\right) \qquad \textit{Equation 1}$$

where $P_{in}$ is the input power, $\eta$ is the mode matching overlap integral (*3*). $T$ is the transmission loss of the mirror coating (10 ppm) and $F$ is the cavity finesse. This expression is valid for all cavity systems influenced by absorption losses (*4*). The remaining path of the splitter was collimated and then focused (Thorlabs, C560TME-B) onto the active area of an APD (Thorlabs, APD430A) enabling collection of reflected resonance light. The reflected signal voltage was sent to a DAQ board (NI, BNC-2120) and monitored using custom software. The transmitted light was collected through the output fiber of the cavity, collimated and focused (Thorlabs, C560TME-B) onto the active area of an APD (Thorlabs, APD430A). The transmitted signal voltage was sent to a diplexer (Mini-Circuits, ZDPLX-2150-S+), the low frequency component (DC-10 MHz) was then directed to the DAQ board (NI, BNC-2120) in order to monitor the transmitted signal. The high-frequency component (50-2150 MHz) was amplified and sent to a frequency mixer (Mini-Circuits, ZP-1MH-S+) in which it was multiplied by the sinusoidal signal of the VCO under a homodyne detection scheme. The resulting frequency components were low-pass filtered (Mini-Circuits, SLP-1.9+) to extract the DC error-signal and directed to the error input of the Pound-Drever-Hall lockbox (PDH, Vescent, D2-125). The locking feedback was supplied to the piezos driving the cavity length via the servo output of the PDH lockbox.

#### Hardware control and data collection software

Experiments were performed using code written in-lab using Python 3 to handle both instrument control and data collection. The code interfaced with a DAQ board (NI, BNC-2120) and oscilloscope (Rigol, DS1104). For DAQ and oscilloscope connections, control was established using the PyLabLib package published under the GPL-3.0 license (doi.org/10.5281/zenodo.7324876). The control code was written together with a graphical user interface (GUI) to streamline user control. The GUI and interactable elements of the program use the Qt framework through PySide2 bindings under the GPLv2 license. A copy of the control software is available for free use under the GPLv3 license at https://github.com/FairhallAlex/ces_tools.

#### Sample preparation

Lyophilized Streptavidin (MilliporeSigma, 189730), Carbonic anhydrase (MP Biomedicals, 0215387910), Aprotinin (MilliporeSigma, A6106) and c-Myc peptide (MilliporeSigma, M2435) were dissolved to 1 $mgml^{-1}$ in phosphate buffered saline (pH 7.4), aliquoted to appropriate volumes and stored at -20 °C. For experimentation, samples were thawed on ice and diluted to the working concentration (0.25 pM-15 pM) in filtered (Ø= 20 nm, Whatman Anotop, WHA68091002) ultrapure Millipore water (18 MΩ, pH 7). To ensure that the measurements would be at the single-molecule level, the optical mode volume was calculated to host an average of 0.7 molecules at the highest working concentration (15 pM).

DNA sequences for each construct were manually designed and validated by NUPACK (www.nupack.org). Commercially available oligonucleotides (Integrated DNA Technologies, Table S 1) were purchased and used without purification. To assemble each structure, corresponding DNA strands (5 μM) were mixed in folding buffer (25 mM HEPES, 100 mM KCl, 10 mM MgCl2, pH 7.4), then annealed in a PCR thermo-cycler (Bio-Rad) by a linear cooling step from 95°C to 20°C over 2 hours (-0.1°C per 10 second). The products were further purified by a size-exclusion chromatography column (Superdex 200 increase 10/300) on a ÄKTA pure system (Cytiva), to remove extra strand and potential aggregates. The final concentration of each construct was determined by Nanodrop (Thermo Fisher Scientific). Samples were stored at 4°C for less than 1 month before use.

**Table S 1.** DNA structures and sequences

| Structure | Sequence |
|---|---|
| 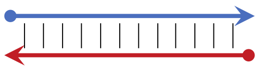 | TGGAGAGAATCGGTCACAGTACAACCG<br>CGGTTGTACTGTGACCGATTCTCTCCA |
| 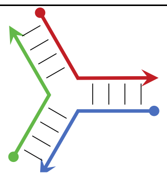 | TGGAGAGAATCGGTCACA<br>GTACAACCGTTCTCTCCA<br>TGTGACCGACGGTTGTAC |

#### Experimental data collection

Single biomolecule diffusion experiments

Prior to introducing proteins, filtered (Ø= 20 nm, Whatman Anotop, WHA68091002) MilliQ water (8 μL) was added to the cavity. The input power in the cavity was controlled with an applied voltage to the VOA (Thorlabs, V600A) to ensure consistent power was used for all comparable experiments. The fundamental transmitted cavity mode was found under active cavity length translation and PDH locked with a proportional gain of -40 dB. The locked position was tuned to the maximum possible transmission using the relative voltage offset on the lockbox (Vescent, D2-125). If spurious events were detected during the lock, the cavity was unlocked and cleaned under a flow of filtered MilliQ water and dried with $N_2$. This process was repeated after protein experiments to ensure the cavity was clean. The transmitted and reflected signals were monitored as a function of time at either 50 kHz or 500 kHz acquisition frequency. Intensity-time traces containing single-molecule events were saved as .csv files every 30 seconds using the custom software described above. Water control experiments were continued until 5 minutes of data in the absence of a spurious signal was collected.

Solutions of proteins (either Streptavidin, Carbonic Anhydrase, Aprotinin or Myc-tag, 0.2-15 pM, 8 μL) or DNA (8 μL, 10 pM) in filtered MilliQ water were introduced into the cavity. The input power in the cavity was controlled with an applied voltage to the VOA to ensure consistent power was used for all comparable experiments. The fundamental cavity mode was found under active cavity length translation and PDH locked with a proportional gain of -40 dB. The transmitted and reflected signals were monitored as a function of time at either 50 kHz or 500 kHz acquisition frequency. Intensity-time traces containing single-molecule events were saved as .csv files every 30 seconds using the custom software described above.

#### Locking bandwidth characterization

The locking bandwidth (LBW) of the cavity was measured by adding a harmonic perturbation ($F_h$) of known frequency and amplitude together with the error signal (e) using a voltage adder to the PI input (Figure S 20). To maintain a linear relationship between the voltage of the error signal during locking and the frequency detuning of the cavity, we optimized the amplitude of the perturbation to 0.2 × the peak-to-peak amplitude of the error signal. While the cavity was locked, we registered the sum of the perturbation signal and the error signal for different frequencies of the perturbation signal. This allowed us to measure the frequency at which the lock ceases to be effective at compensating for the perturbation to the system to determine the 0 dB gain value and thus the LBW of ~5 kHz (Figure S 10, Figure 5A).

#### Noise background experiments

We measured the noise profile of the locked cavity from the error signal data. The voltage amplitude of the error signal was converted to its corresponding cavity frequency shift using the scanned error signal slope calibrated with the modulation sidebands at 2.6 GHz. From the calculated Fourier transform of the error signal, we extracted the RMS values of the cavity resonance frequency shift. We performed this measurement for different proportional gain settings where the maximum noise suppression was reached for the maximum locking bandwidth at ~5 KHz. The noise floor was measured from the off-resonance error signal. This value, as well as the cavity finesse and gain determine the LBW. We found effective noise suppression of external perturbations by the PDH locking loop at the level of the noise floor throughout the LBW. While the source of low frequency noise is mainly due to ambient acoustic waves, mechanical waves that couple through the cavity system via contact points to the optical table, higher frequency noise (above the LBW) inherent to the electronic control systems is suppressed using a low pass passive filter before the piezo connection. Mechanical resonances of the cavity are expected to appear at frequencies above the LBW, however its magnitude is below the detector noise floor (*2*), at the level of $10^3$ $HzHz^{-1/2}$ (Figure S 14). The high mechanical passive stability of the cavity and the active PDH defines a frequency region where the cavity can be more sensitive to internal perturbations as described in the main text.

#### Voltage pulse experiments

To demonstrate the mechanism described in the main text (Figure 5C), we induced controlled perturbations to the locked cavity to mimic molecular perturbations to the cavity. The passage of molecules into the cavity results in an increase in the average refractive index, and a consequent decrease in frequency.  This frequency decrease was mimicked by transiently, slightly increasing the length of the cavity by applying a voltage pulse to the piezo. Based on the measured duration of molecular transit events (Figure S 5) and their prominence, we initially selected the parameters of the pulse, including duration and amplitude, to approximately induce similar detuning magnitudes and cavity transmission profiles (Figure S 19). Square pulse signals of 140 µs-1 ms duration (Figure S 19) and 1 Hz repetition rate were produced to affect the cavity length while the cavity was locked. The pulse produced by a function generator (Keysight, DSOX1204G) was added to the servo output of the lockbox; this combined signal was then directed to the piezos. The transmission signal of the locked cavity over time was recorded to ensure that all perturbation events and pulse signals were correlated (Figure S 19A). A small reaction delay was seen due to the length of the cables that drive the piezo, the reaction time of the electromechanics and the transmission signal to the DAQ card. The transmission profile (Figure S 19, blue trace) is the result of the step down voltage perturbation (gray) which produced a steep reduction of the locked transmission signal due to the photo-thermal effect (a), this was followed by a brief recovery to the locked state by the PI feedback (b) followed by a second descent of the transmission signal as the step-up voltage of the pulse shifts the cavity in the opposite direction (c), finally the PI control recovers the locked state (d). When the pulse is generated, the resulting perturbation has a profile that resembles the transmitted signal profiles of the cavity due to the molecular interaction, i.e. a shift to smaller frequencies (Figure 2A, Figure S 5).

#### Photothermal broadening and thermo-optic coefficient

Optical microcavities with small mode volumes are susceptible to dynamic photothermal non-linearities that originate from the build-up of intense optical fields causing the temperature to increase in the cavity. Coupling between the heat and cavity resonance frequency can result in drift of resonance frequencies and cooling cascades (*5*).

These photothermal dynamics depend on the thermo-optic coefficient (dn/dT) of the medium dominating the thermal dissipation. Increasing the temperature of a medium with a negative dn/dT will result in a decrease in the refractive index, and a consequently lower cavity resonance wavelength (high frequency). When the cavity length or laser frequency is scanned, the resonance may drift to higher or lower wavelengths depending on the direction of the scan and the cavity-laser detuning. When the cavity length is scanned to longer lengths, the optimal resonance condition (Equation 2),

$$m\lambda = 2nL \qquad \textit{Equation 2}$$

where *m* is an integer, *λ* is the wavelength, *n* is the refractive index of the medium and *L* is the cavity length, appears to change as n decreases, resulting in a need for still higher lengths, and resulting in a distorted line shape, (Figure S 17).  This behavior is expected in FFPCs when the medium is air and the thermal dissipation is dominated by the mirror coatings, where an “effective” negative dn/dT (*6*) due to thermal expansion results in a shorter cavity length (*3*). A negative dn/dT is also expected when the medium is water (*7*). We characterized the photothermal behavior in our FFPCs with water, by actively scanning the cavity to increasing length and observing the direction of photothermal broadening in water (Figure S 17). A positive voltage gradient was applied to one of the piezos and confirmation that the positive voltage corresponded to increasing cavity length was confirmed by tuning the wavelength of the pump laser and observing a shift of the resonance position to higher voltages with a lower pump wavelength. The direction of the broadened resonance was observed to be towards longer cavity lengths, thus lower wavelengths, as expected for a system dominated by a medium with a negative thermo-optic coefficient.

#### Photothermal bandwidth measurement

Photothermal broadening of cavity lineshapes as a function of applied ramp speed were recorded in order to quantify the photothermal bandwidth of the cavity in water (Figure S 18A). Resonances were recorded in transmission as a function of time and the cavity length was tuned using a ramp signal applied to the piezos from a function generator (Keysight, DSOX1204G).  Phase modulated sidebands at a known frequency (2.6 GHz) were applied to be used as a frequency calibration marker to each trace. Initially, the ramp frequency and amplitude were chosen to minimize the photothermal effects in the cavity. This was determined by matching resonance lineshapes during both the increase and decrease of ramp voltage. This trace was used to measure the linewidth of the cavity. From there, both the ramp speed was scanned and traces were recorded for both the increase and decrease in ramp voltage components of the ramp signal.

#### Data analysis

Single-molecule diffusion and autocorrelation analyses

Analysis of transmitted and reflected signals resulting from perturbation by single biomolecules was performed with custom written code using Python 3. Biomolecule and water control data collected at a single input power were first normalized to the maximum (for reflection) or minimum (for transmission) signal intensity to enable all signal peaks at a comparable power to be selected with a single threshold between 0-1. Single events were identified and analyzed using the SciPy.signal find_peaks package. Signal peaks were selected at a prominence threshold of 0.35. A temporal filter of 2 ms was applied to ensure that only peaks separated in time by greater than 2 ms were selected. Temporal widths were determined at the full width at half maximum of the event, and prominences were determined as the vertical distance between the maximum of the peak and the local background intensity.

Autocorrelation analysis was performed to quantify the temporal behavior of the single-molecule events in the intensity-time traces with custom code written in Python 3. A normalized function was generated by comparing time-shifted values to one another (Equation 3):

$$G_k = \frac{\sum_{i=1}^{N-k}(Y_i - \bar{Y})(Y_{i+k} - \bar{Y})}{\sum_{i=1}^{N-k}(Y_i - \bar{Y})^2} \qquad \textit{Equation 3}$$

where $k$ is the number of time steps, $N$ is the total number of points, $Y_i$ is the intensity for a specific time and $\bar{Y}$ is the average intensity. Several 30 s intensity-time traces for proteins diffusing in the locked cavity were concatenated. To ensure that the background was approximately continuous between files, the minimum of each file was found and then subtracted from every point. Events were identified with an intensity threshold 2.5 standard deviations from the mean and used to generate an autocorrelation trace. The function sm.tsa.acf from the package statsmodels.api, was used to generate the autocorrelation function. These functions were plotted for the four different proteins (streptavidin, carbonic anhydrase, aprotinin and Myc-tag, Figure 3A), and the values for different decay times were extracted. The time to decay to 40% of the autocorrelation time is shown to be linear with protein radius, Figure 3B. This linear trend was preserved for a large range of decay values (Figure S 11), demonstrating the robustness of the analysis. A copy of the analysis software is available for free use under the GPLv3 license at https://github.com/FairhallAlex/ces_tools.

#### Locking bandwidth analysis

To extract the effective LBW of the system, we computed the Fourier transform of the signal measured at point S for each perturbation frequency (Figure S 20). The amplitude at the perturbation frequency were extracted and normalized with respect to the input signal $F_h$ and converted to decibels. The result of the normalized values are shown as a function of input frequency of the sine wave (Figure 5A, Figure S 10). The corresponding LBW was determined to be ~ 5 kHz at the 0 db crossing point. At larger frequencies the signal was amplified due to a change in phase of the control circuit known as the servo bump. At even higher frequencies the lock had no influence on the perturbation signal and the relative intensity remains at 0 db.

#### Signal-to-noise ratio determination

The data traces were smoothed using various bin sizes to generate a moving average, following:

$$F_{smoothed,i} = \frac{\sum_{j=i}^{i+(n-1)} F_{raw,j}}{n} \qquad \textit{Equation 4}$$

where $n$ is the bin size and $F_i$ is the value of the function at index $i$. This smoothing procedure was performed for a series of different bin sizes from $n$=1 to $n$=250. Following smoothing, the trace was normalized. Smoothing of the data was only performed for SNR calculations. To then calculate the signal-to-noise ratio (SNR), the standard deviation of the background ($\sigma_N$) was taken from the longest segment of background in the trace. The signal

(*S*) was then calculated from the amplitude of the peaks and the SNRs for the proteins measured in this paper (Figure S 6) were calculated by:

$$SNR = \frac{S}{\sigma_N} \qquad \textit{Equation 5}$$

quantifying the SNR for each peak under each smoothing bin size. Giving the maximum average SNR, a bin size of 190 which was used for all datasets.

For comparison with other techniques, where provided, SNR values were taken directly from references (*8–12*). For reference (*13*), data was extracted into a .csv from Figure 4a using WebPlotDigitizer. The SNR was then calculated by taking the maximum of the signal and dividing it by the standard deviation of a region without signal. For reference (*14*)**,** the resonance shift was taken to be 5 am as a high estimate from Figure 5e and divided by the reported noise level of 9.6 × $10^{-4}$ fm. For reference (*15*), the standard deviation of the noise was calculated by estimating 3σ from Figure 5b and dividing that by 3. The SNR was then calculated using the mean reported value of 2.5 fm for the 8-mer and then dividing by the previously calculated standard deviation.

#### Photothermal bandwidth determination

To determine the photothermal bandwidth, the frequency shift of the photothermally broadened peak was calculated for each corresponding ramp speed. The photothermally broadened peak for each ramp speed had the larger distance between the left sideband and the main peak, which was determined to be the interpeak distance. Using this information, the frequency shift for each ramp speed was then calculated by calculating the difference between the photothermally broadened interpeak distance and the photothermally narrowed interpeak distance. The frequency shift as a function of the ramp speed (Figure S 17) were then fitted to Equation 6 (*6*).

$$\beta(x) = \frac{\beta_{ad}}{1 + x\tau} \qquad \textit{Equation 6}$$

where $\beta_{ad}$ and $\tau$ corresponded to the adiabatic thermal resonance shift and the thermal reaction time constant respectively. $\beta(x)$ is the frequency shift and *x* is the linewidth. $\beta_{ad}$ was calculated to be 8.28 linewidths and $\tau$ was calculated to be 23.7 linewidths per microsecond. Then using Equation 7:

$$f_{th} = \frac{1}{2\tau}$$ *Equation 7*

$\tau$ was converted to $f_{th}$ (21 kHz) which is the bandwidth for photothermal resonator length stabilization.

#### Simulations and calculations

##### Calculated resonance shift

We can use analytical expressions to determine frequency shifts from small objects. Essentially, these are the shifts without any additional photothermal enhancements, and can be thought of as the observed shifts in cavities with extremely small circulating power. We used two approaches, first we applied Equation 8 following the approach by Kohler *et al* (*16*) (Figure S 13B). Here, the polarizability of the molecule is calculated via the Lorentz-Lorenz equation, Equation 9, for mixtures weighted by the overlap between the mode volume and the molecule and divided by the volume of the molecule:

$$\langle\alpha\rangle = \frac{\alpha}{V_{mol}}\int_{V_{mol}} dV_{np}\left(\frac{\omega_0}{\omega(z)}\right)\exp\frac{-2(x^2+y^2)}{\omega(z)^2}\cos^2(kz)$$ *Equation 8*

where

$$\alpha = 4\pi r^3\epsilon_0\frac{n_{mol}^2 - n_{water}^2}{n_{mol}^2 + n_{water}^2}$$ *Equation 9*

$$\omega(z) = \frac{\omega_0}{\sqrt{1+\left(\frac{z}{z_0}\right)^2}}$$ *Equation 10*

The mode area and the Rayleigh length are:

$$\omega_0^2 = \frac{L\lambda_m}{\pi}\sqrt{\frac{G_1G_2(1-G_1G_2)}{(G_1+G_2-2G_1G_2)^2}}$$ *Equation 11*

$$z_0 = \pi\omega_0^2/\lambda_m$$ *Equation 12*

correspondingly. Here $L$ is the cavity length, $\lambda_m = \lambda_0/n_{water}$ is the wavelength in the medium, and the cavity geometry parameters are $G_i = 1 - L/R_i$ , where $R_i$ are the radius of curvature of each mirror. The cavity frequency shift is:

$$\Delta\nu = \frac{\langle\alpha\rangle c}{2\lambda_m\epsilon_0 V_{mode}}$$ *Equation 13*

where $c$ is the speed of light in vacuum and the mode volume is $V_{mode} = \pi\omega_0^2 L/4$. This approach differs from Kohler *et al* (*16*) as the molecules are significantly smaller than the mode volume and therefore the mode shape shifts a negligible amount on the length scale of the molecule.

For the four proteins interrogated in this work the corresponding resonance shifts ($\Delta\nu$) and losses that would be induced by interaction with the cavity mode were calculated (Table S 2).

**Table S 2.** Calculated resonance shifts induced by proteins streptavidin, carbonic anhydrase, aprotinin and Myc-tag upon interaction with the mode volume of the FP microcavity. This approach was taken from Kohler *et al* (*16*).

| Protein | Refractive index | Protein radius (nm) | $\Delta\nu$ (kHz) |
|---|---|---|---|
| Streptavidin | 1.43 | 2.80 | 49.0 |
| Carbonic anhydrase | 1.43 | 2.10 | 21.0 |
| Aprotinin | 1.43 | 1.45 | 6.7 |
| Myc-tag | 1.43 | 0.75 | 0.96 |

The second approach was adapted from Su *et al* (*14*). First, the polarizability was calculated as previously shown using Equation 8. The wavelength shift was then calculated following Equation 14:

$$\Delta\lambda_{max} = \frac{\langle\alpha\rangle\left[\frac{E_0^2(r_e)}{E_{max}^2}\right]}{2V_m}\lambda \qquad \textit{Equation 14}$$

where *r* was the particle radius, $\lambda$ the free space wavelength, $\frac{E_0^2(r_e)}{E_{max}^2}$ was calculated to be 1/5.5 for the microtoroid (*14*) and 1/4.8 for the FP microcavity, since the molecule is able to overlap with the mode maximum. To calculate the resonance shift expected from interaction of the proteins with a toroidal microresonator the mode volume, $V_m$, was taken from Su *et al* (*14*) and was equal to 330 μm$^3$. In order to compare to the expected resonance shift from the FP cavities used in this work, $V_m$ was calculated to be 80 μm$^3$. Given these values, the resulting frequency shifts were calculated as shown in Table S 3.

**Table S 3.** Calculated resonance shifts induced by proteins streptavidin, carbonic anhydrase, aprotinin and Myc-tag upon interaction with the optical mode of a toroidal microcavity. This approach compares the calculated shifts given the mode volume of a toroidal microcavity and our FP microcavity and was taken from Su *et al* (*14*).

| **Protein** | **Refractive index** | **Protein Radius (nm)** | **$\Delta\nu$ toroid (kHz)** | **$\Delta\nu$ FP (kHz)** |
|---|---|---|---|---|
| Streptavidin | 1.43 | 2.8 | 3.796 | 15.064 |
| Carbonic anhydrase | 1.43 | 2.1 | 1.601 | 6.355 |
| Aprotinin | 1.43 | 1.45 | 0.530 | 2.092 |
| Myc-tag | 1.43 | 0.757 | 0.075 | 0.297 |

The magnitude of the resonance shifts induced from the interaction between the proteins and the mode of a toroidal microresonator are less than the shifts from the same interaction in our FP cavities. The confinement of the optical mode in the medium outside of the dielectric material in FP microcavities allows for stronger overlap between the molecule and the mode when compared to a toroid, in which the mode is confined within the dielectric material (Table S 2). Furthermore, the smaller mode volume in the FP microcavity facilitates stronger light-matter interactions further resulting in larger resonance shifts compared to those achieved with a toroidal microcavity (Table S 3).

#### Molecular velocity distributions and molecular mean-square-displacement power-spectral-density

The velocity distribution of particles undergoing free Brownian motion in solution follow Equation 15 and Equation 16 (*17*):

$$m^* = m_p + \frac{1}{2} m_f \qquad \textit{Equation 15}$$

$$f(v) = \sqrt{\frac{m^*}{2\pi k_B T}} e^{-\frac{m^* v^2}{2k_B T}} \qquad \textit{Equation 16}$$

for particle of mass $m_p$, solution mass $m_f$, and solution temperature $T$. An effective mass was calculated in order to include the effects of the solution on the acceleration of the particle. With these equations, one can approximate the likelihood of a particle to move with a given velocity (Figure S 15A). The smaller the particle, the wider the velocity distribution becomes.

The mean squared displacement power spectral density (MSDPSD) of a particle undergoing free Brownian motion can be calculated with the Equation 20 (*17*) , derived as a solution to the Langevin equation:

$$D = \frac{k_B T}{6\pi\eta r} \qquad \textit{Equation 17}$$

$$\tau_f = \frac{r^2 \rho_f}{\eta}, \tau_p = \frac{m}{6\pi\eta r} \qquad \textit{Equation 18}$$

$$\phi_f = \frac{1}{2\pi\tau_f}, \phi_p = \frac{1}{2\pi\tau_p} \qquad \textit{Equation 19}$$

$$P_{free}(f) = \frac{D}{\pi^2 f^2} \frac{1 + \sqrt{f/2\phi_f}}{(\sqrt{f/2\phi_f} + f/\phi_p + f/9\phi_f)^2 + (1 + \sqrt{f/2\phi_f})^2} \qquad \textit{Equation 20}$$

for particle radius $r$, particle mass $m$, solution density $\rho_f$, solution viscosity $\eta$, and solution temperature $T$. The $\tau$ terms are time constants, $\tau_f$ related to inertia of the surrounding fluid, and $\tau_p$ to the inertia of the particle itself. We note that this version of the equation is for a free particle (not confined by an optical trap). This equation was plotted for the four proteins using known mass values and radius values for streptavidin, carbonic anhydrase, and aprotinin, with a calculated radius for Myc-tag (Figure 5A, Figure S 15B). A range from 1 kHz to 1 MHz was used, and the integral over the range 5 kHz to 21 KHz was calculated, giving 0.00875 μm$^2$ for Myc-tag, 0.00457 μm$^2$ for aprotinin, 0.00315 μm$^2$ for carbonic anhydrase, and 0.00237 μm$^2$ for streptavidin.

Simulated photothermal bandwidth determination

Finite element simulations were performed using COMSOL (version 6.0). The adiabatic model (*5*) was used to solve for the stable equilibrium of the cavity resonance frequency/length/wavelength under high circulating power conditions. While the cavity is locked, the heat generated by the absorbed circulating power is dissipated by the heat conduction into the surrounding medium. The cavity equilibrium resonance frequency is determined by the heat conductance of the system (K) (assuming no convection or radiation). The thermal conductance of the system was calculated by considering a volume of water, equivalent to the optical mode volume, which is heated according to Equation 22. If $a_{water}$ is the absorbed power in water:

$$a_{water} = \pi \frac{1}{F_{air}} - \frac{1}{F_{water}} \qquad \textit{Equation 21}$$

where F is the finesse in air/water respectively, the total absorbed power is:

$$P_{abs} = P_{circ} a \qquad \textit{Equation 22}$$

here, $P_{circ}$ is defined in Equation 1 as the circulating cavity power. Consequently, the total power flow across a gaussian surface surrounding the mode volume of the cavity is measured on a 200 μs time-scale. This integration time was arbitrarily chosen to be much longer than the measured photothermal time-constant of the system, determined to be 7.57 μs. Based on the distribution of the optical mode (Figure S 16) for the geometry of cavity one, a cylinder of water with radius 1.25 μm and length 19 μm was used as the heat source to the model. A volume of water defined by a sphere (radius 25 μm) acted as a heat sink for the system. A glass cylinder of diameter 125 μm was placed tangent to each circular face of the heat source, representing the fibers. Using the Heat Transfer in Solids and Fluids module, a constant power density of 42.2 GW/m$^3$ was applied to the cylinder. This power density was calculated based on the circulating power of the cavity and the absorption of water, giving 3.93599 μW of absorbed power over the volume of the cylinder. The system was set to an initial temperature of 293.15 K, and was allowed to evolve with this power input, approaching a temperature of 293.28 K.

To estimate the thermal relaxation time, we initialized the system at the equilibrium temperature and studied its characteristic temperature decay profile. The resulting temperature over time data was then fit with an exponential decay curve (Figure S 18B). This yielded a time constant of 7.57 μs, consistent with a thermal bandwidth of 66.05 kHz approximately 3-fold larger than the measured value of 21 kHz (Figure S 18A). This small discrepancy is due to non-idealities not considered in the simulation, such as additional contact points with the ferrule which were not considered.

#### Fiber Cavity Mechanics

Finite element simulations on mechanical fluctuations of the cavity were performed using COMSOL (version 6.0). The fiber cavity was modelled (Figure S 22) including the fibers (glass), ferrule (glass) and piezos (Lead zirconate titanate with Young's Modulus 82.1 GPa and Poisson Ratio 0.39) (*18*), and glass plate (Figure S 22). The piezos (2.5 mm × 2.3 mm × 2.5 mm) sat on top of the glass block (20 mm × 7 mm × 3 mm). The slotted ferrule section has the left half (3.4 mm × 1.25 mm × 1.25 mm) separated from the right (4.6 mm × 1.25 mm × 1.25 mm) by a gap of 125 μm. The bore was placed 0.833 mm from the bottom of the ferrule, with a radius of 65.5 μm. There was a half cut through the right ferrule, 0.5 mm from the full gap between left and right. The fibers were each placed within the bore, tangent to the bottom, with a radius of 125 μm. A 19 μm separation between the fibers defined the optical cavity. The water was modeled as an ellipsoid (1 mm × 0.6 mm × 1.25 mm), centered on the plane of the top of the ferrule, directly above the gap between the fibers. The parts of this ellipsoid clipping with the fibers and ferrule were removed. The two modules used for this simulation were Solid Mechanics for all the glass components and piezos, and Pressure Acoustics, Frequency Domain for the water components. Multiphysics boundaries between the two were included. The eigenfrequencies and eigenmodes of this system both with and without water were calculated. Eigenmodes appearing in both the air and water simulations were used for further calculations.

The noise spectral density was calculated as previously described (*2*). From the simulations, the effective masses of the eigenmodes are first calculated, following the equation:

$$m_{eff} = \frac{\int_V dV \rho(x,y,z) \cdot |\boldsymbol{u}(x,y,z)|^2}{max_V(|\boldsymbol{u}(x,y,z)|)^2} \qquad \textit{Equation 23}$$

where V is the volume of the simulation, $\rho(x,y,z)$ is the density at a given position, $\boldsymbol{u}(x,y,z)$ is the displacement field at a given position, and $max_V$ is the maximum within the volume of the simulation. The zero-point motion of the modes is then calculated, following the equation:

$$x_{ZPM} = \sqrt{\frac{\hbar}{2m_{eff}\Omega_m}} \qquad \textit{Equation 24}$$

where $\Omega_m$ is the angular frequency of eigenmode m. The optomechanical coupling rates are then calculated, following:

$$G = \frac{-2\pi\nu_0}{L_{cavity}} \qquad \textit{Equation 25}$$

$$g_0 = G \cdot x_{ZPM} \cdot \left(\frac{u_{x,mirror\ 1}}{max_V(|\boldsymbol{u}(x,y,z)|)} - \frac{u_{x,mirror\ 2}}{max_V(|\boldsymbol{u}(x,y,z)|)}\right) \qquad \textit{Equation 26}$$

where $L_{cavity}$ is the length of the cavity (here 19 μm), $u_x$ is the maximum displacement in the x direction for an arbitrary perturbation (parallel to the cavity optical axis), and $\nu_0$ is the optical frequency being coupled to.

A linewidth of 1000 Hz ($\Gamma m$= 6283.185) was used as an approximation for all of the modes, as done in Saavedra *et al* (*2*). With these linewidths, the frequency noise spectral density can be calculated, following the equation:

$$S_v(f)^2 \approx \frac{2{g_0}^2}{4\pi^2} \cdot \frac{2\Omega_m}{\hbar} \cdot \frac{2\Gamma_m k_B T}{\left(\Omega^2 - {\Omega_m}^2\right)^2 + {\Gamma_m}^2\Omega^2} \qquad \textit{Equation 27}$$

where $\Omega$ is the noise angular frequency ($\Omega = 2\pi f$), temperature is T, and Boltzmann constant is $k_B$.

The noise floor of the cavity is an order of magnitude lower than the detector noise floor (Figure S 14) within the molecular velocity bandwidth. A stepwise integral of $S^2$ from the locking bandwidth (5 kHz) to the photothermal bandwidth (21 kHz) was calculated, and then the square root taken, giving the integrated noise within the selected region. The integrated noise over the observation bandwidth is on the order of the resonance shift expected from the smallest molecule, Myc-tag (Table S 2).

### Supplementary Figures and Tables

**Table S 4.** Parameters of the cavities used in this work. The cavities used to collect the data shown in the figures of the main text are indicated in this table, the cavities used to collect the data shown in the supplementary figures are indicated in the respective figure legends.

| Parameter | Cavity one | Cavity two | Cavity three | Cavity four |
|---|---|---|---|---|
| $\lambda_{pump}$ (nm) | 660 | 660 | 660 | 760 |
| Finesse | 37450 | 17909 | 21780 | 30000 |
| Cavity length (µm) | 19 | 19 | 24 | 20 |
| Δv (MHz) | 206.87 | 398.83 | 286.9 | 261 |
| ROC mirror 1 (µm) | 122.4 | 116.9 | 60.9 | ~170 µm |
| ROC mirror 2 (µm) | 97.7 | 105.5 | 67.5 | ~170 µm |
| Figure 1 | X | | | |
| Figure 2 | X | | | |
| Figure 3 | X | | | |
| Figure 4 | | X | | |
| Figure 5 | | | | X |

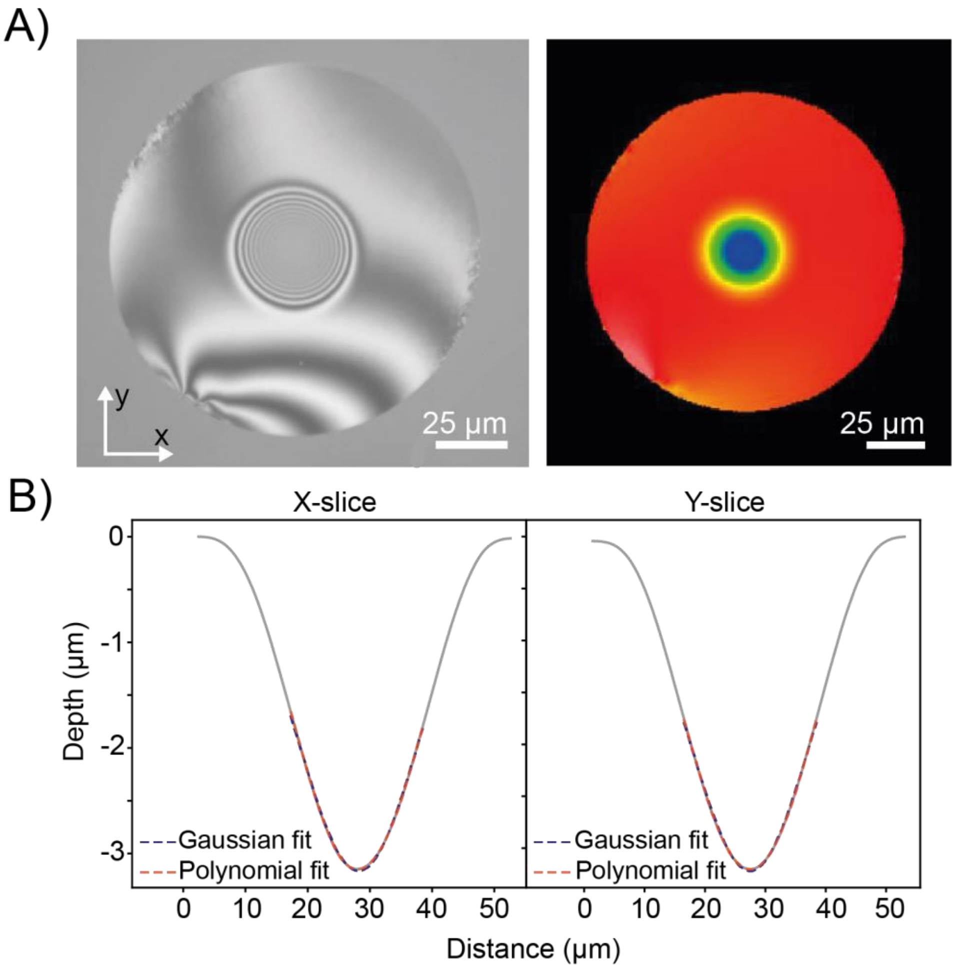

**Figure S 1. A)** Interferometry profiles of the concave depression created by $CO_2$ laser ablation into the face of an optical fiber. **B)** Resulting 2D depth profiles created from X and Y slices of the interferometry images, these are fitted with appropriate functions to define the radius of curvature (ROC) and diameter of the ablation.

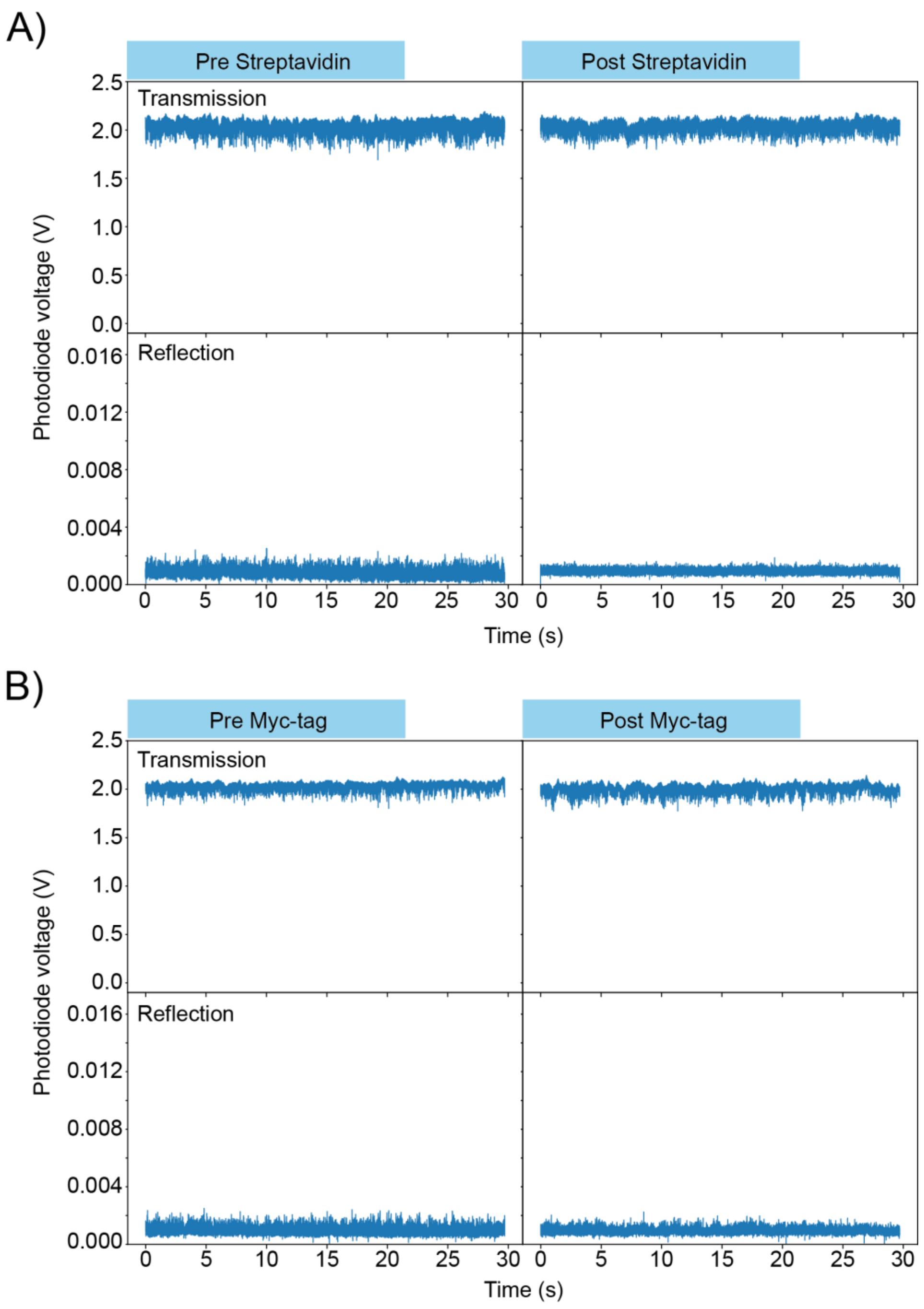

**Figure S 2.** Representative intensity vs time traces of a locked water filled cavity in the absence of molecules in both transmission and reflection either **A)** pre and post introduction of streptavidin or **B)** pre and post introduction of Myc-tag. The lack of signal here demonstrates that the transient perturbations (Fig 2A) orignate from molecules. Data collected in cavity one.

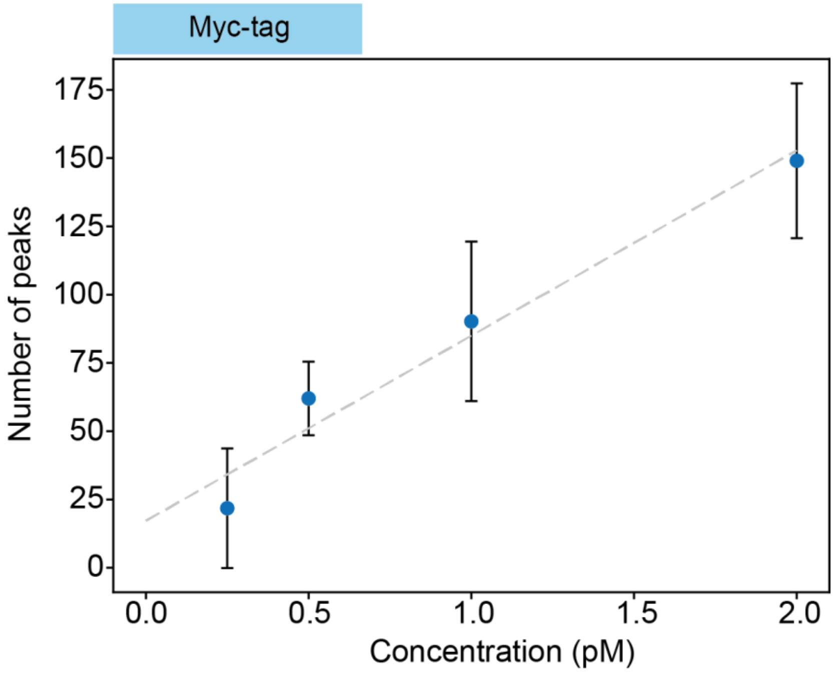

**Figure S 3.** Number of detected peaks induced by Myc-tag as a function of the protein concentration. Error bars represent the standard deviation across datasets. This scaling demonstrates that the peaks are induced by interactions between biomolecules and the cavity-mode. Data collected in cavity two.

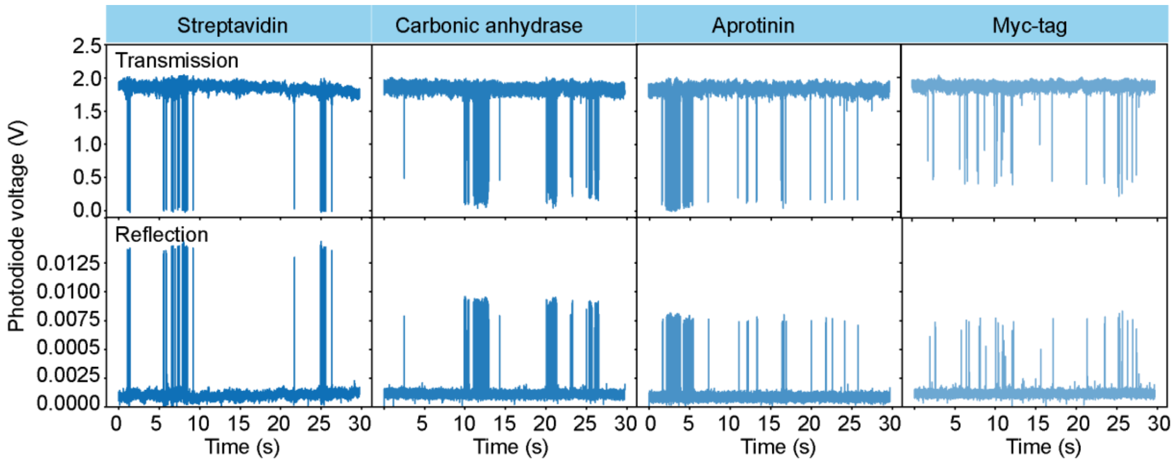

**Figure S 4.** Full 30 s time traces of protein data displayed in Figure 2A in the Main Text. Data collected in cavity one.

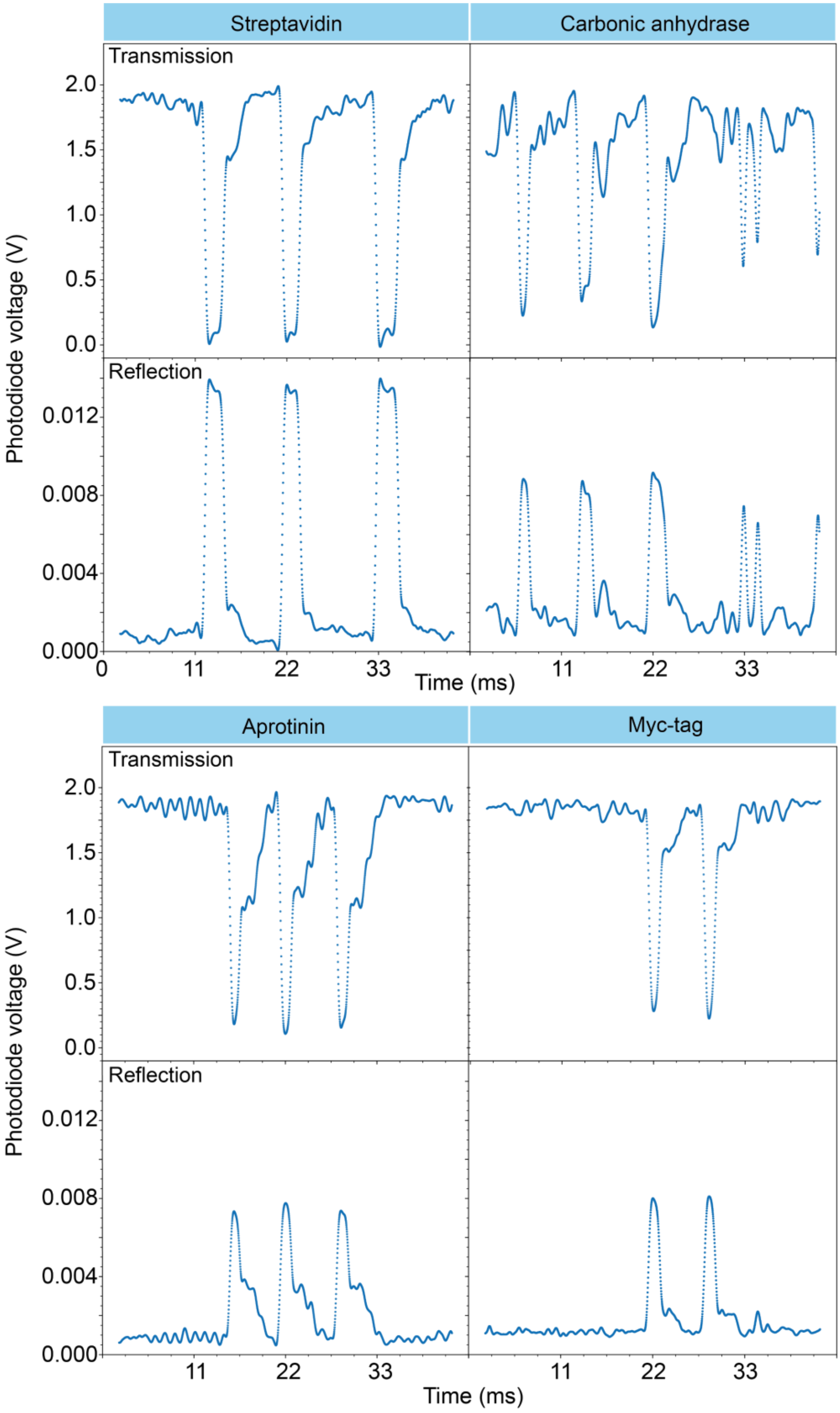

**Figure S 5.** Representative 44 ms traces of proteins Streptavidin, Carbonic Anhydrase, Aprotinin and Myc-tag perturbing the cavity mode in both transmission and reflection. Data collected in cavity one.

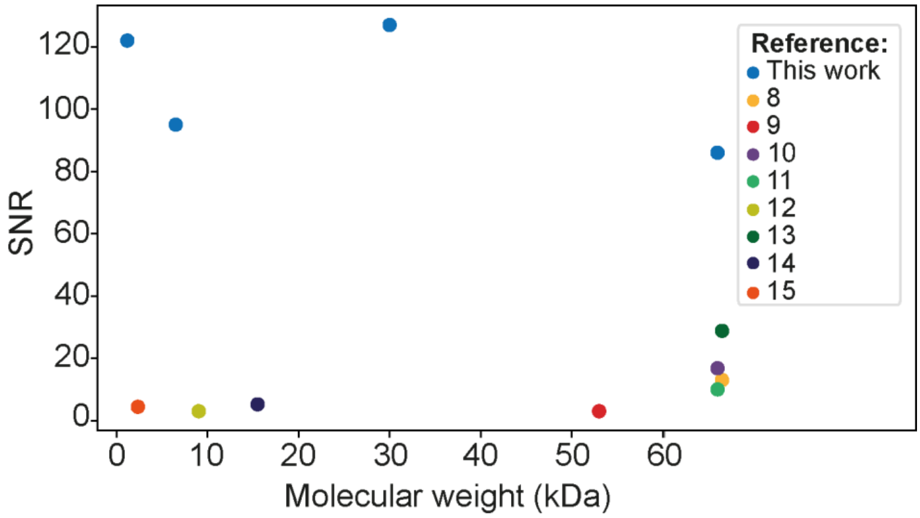

**Figure S 6.** Comparison between signal to noise ratios of this work and other label-free, single-molecule studies (*8–15*). Single-molecule diffusion data shown in Fig 2A and collected in cavity one.

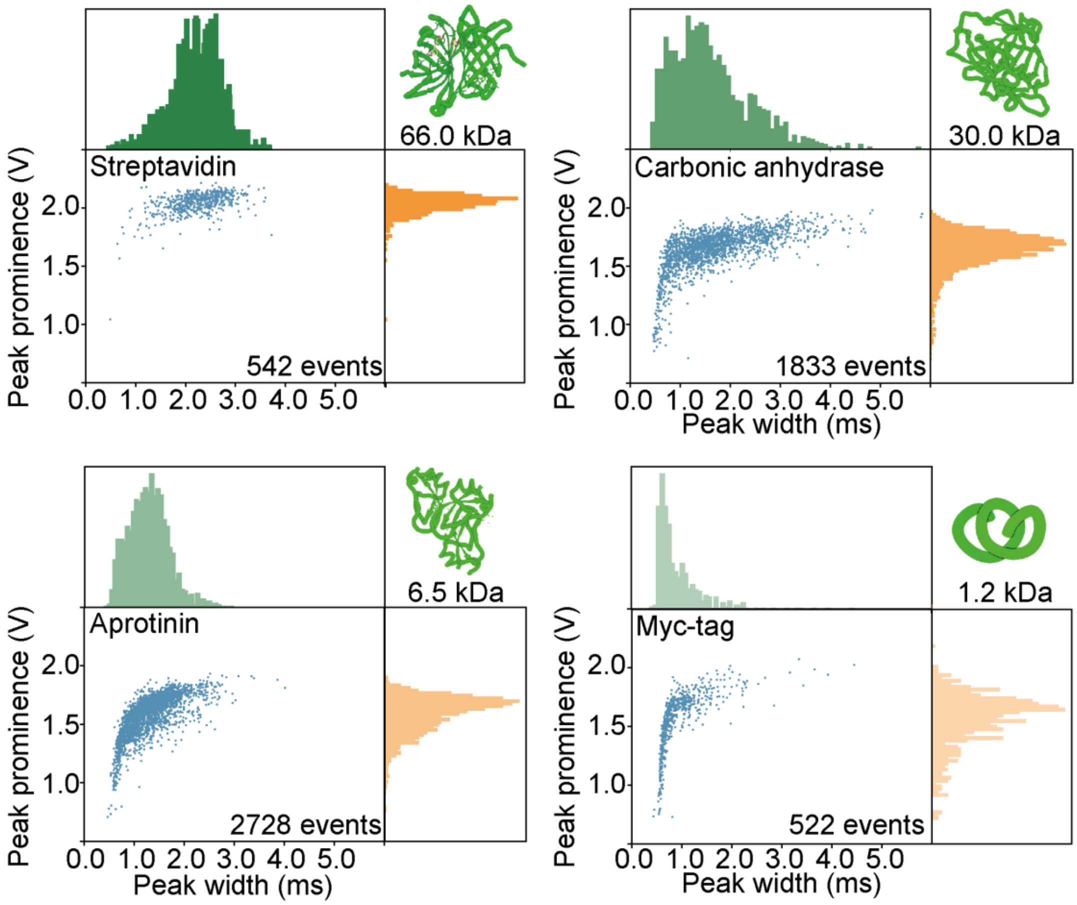

**Figure S 7.** 2D plots and accompanied histograms of the extracted prominences and temporal widths of the transmitted signals. The reflection signals are displayed in Figure 2A in the main text. Data collected in cavity one.

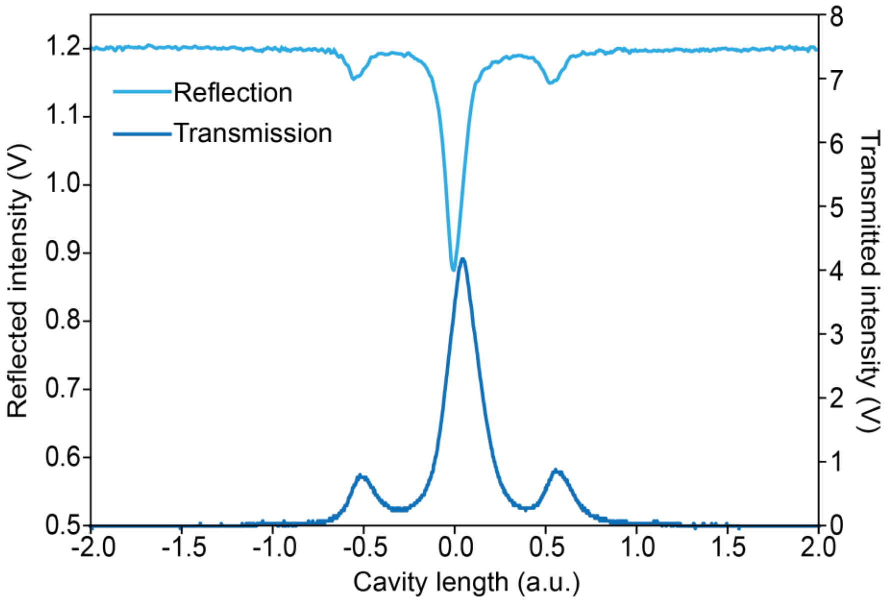

**Figure S 8.** Example resonances in reflection and transmission under cavity length translation. The frequency of the reflected signal is offset relative to the transmitted signal by 309 MHz due to mode matching leading to dispersion. Data collected with cavity one.

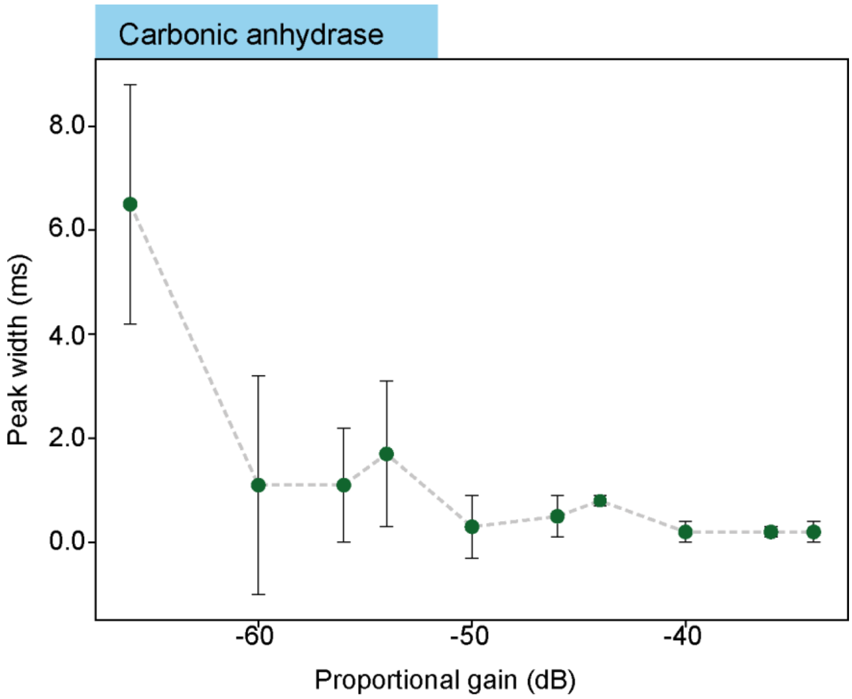

**Figure S 9.** Temporal widths of carbonic anhydrase diffusion events as a function of the locking bandwidth of the PDH determined by the proportional gain of the PI control. This relationship can be explained by the inverse relationship between the molecular velocity filter bandwidth and the proportional gain values. As the gain is increased the velocity filter bandwidth narrows, resulting in detection of a distribution of faster moving molecules with narrower peak widths. Data was collected at proportional gain settings >-50 db where the mean temporal width was no longer influenced by the locking bandwidth. Error bars represent the standard deviation of temporal widths across all analyzed peaks. Data collected with cavity three.

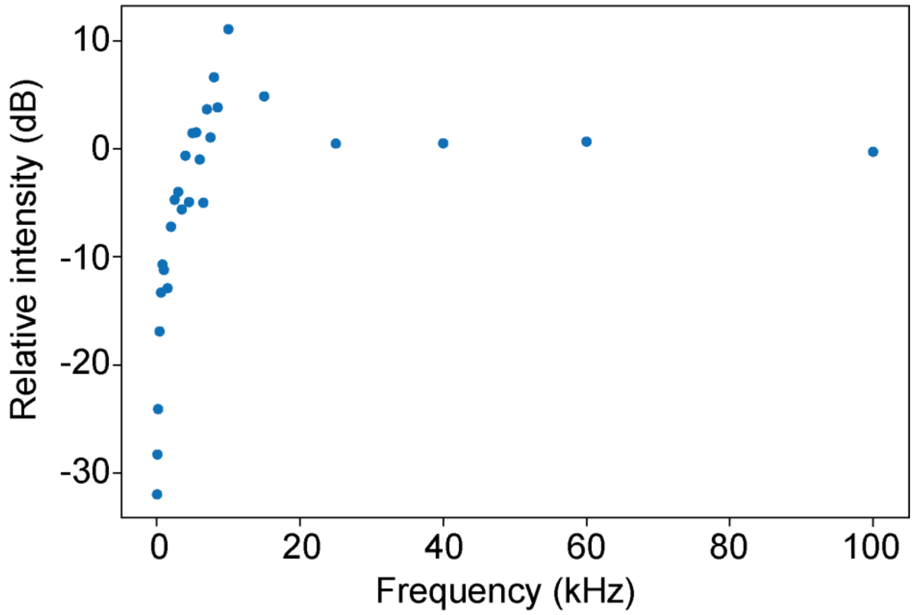

**Figure S 10.** Determination of the locking bandwidth. The frequency of the locking bandwidth is detemined at the relative intensity crossing at 0 dB and is approximately 5 kHz. Data collected with cavity two.

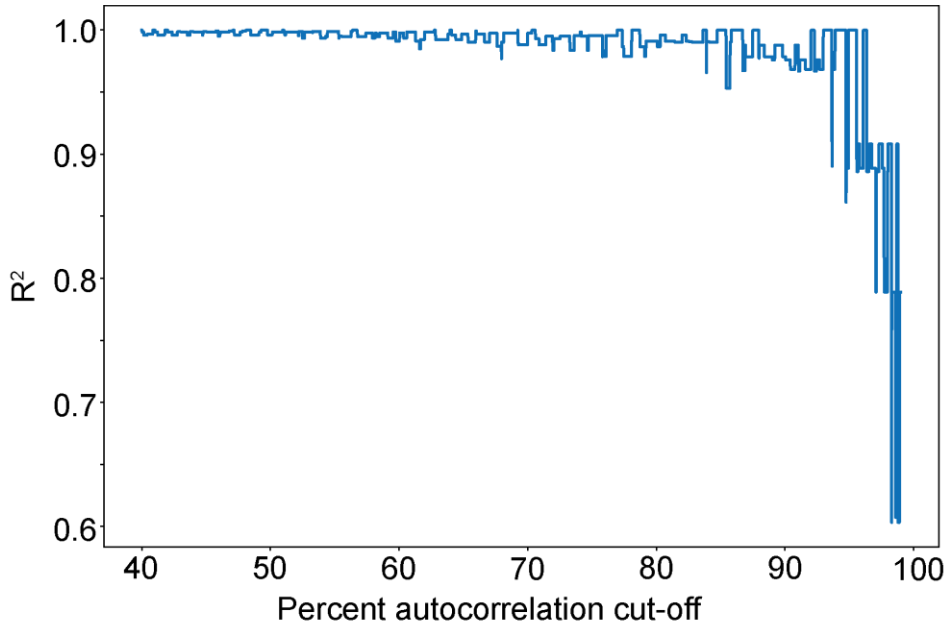

**Figure S 11.** Linearity of the autocorrelation function as a function of protein radius measured via the $R^2$ of the linear fit versus the percentage autocorrelation cut-off shown in Figure 3 in the main text. This linear trend was clearly preserved for a large range of decay values, demonstrating the robustness of the analysis.

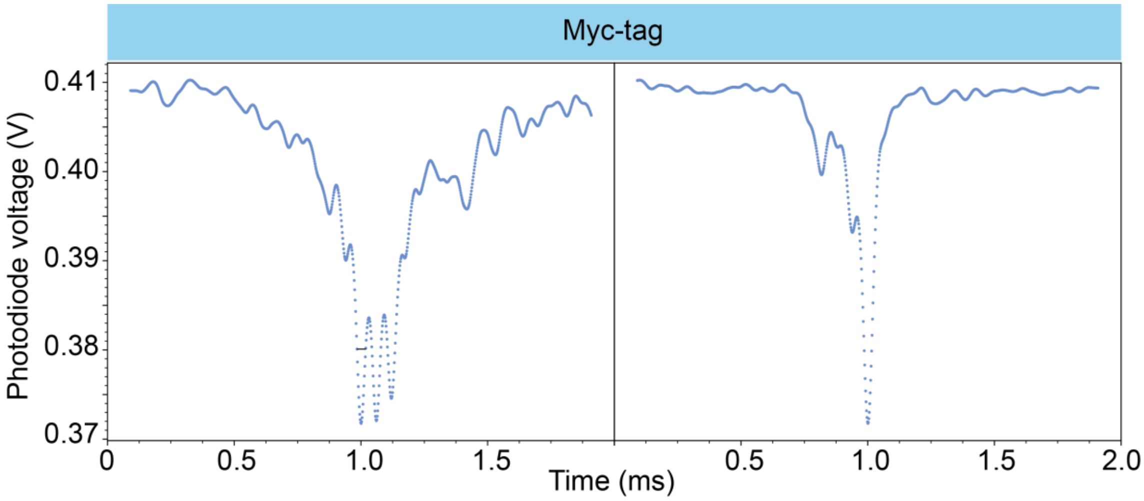

**Figure S 12.** Single diffusion events of <1 nm protein, Myc-tag, are resolvable with 2 μs temporal resolution. Representative single Myc-tag events collected with 500 kHz acquisition frequency. Data collected using cavity two at a concentration of 1 pM.

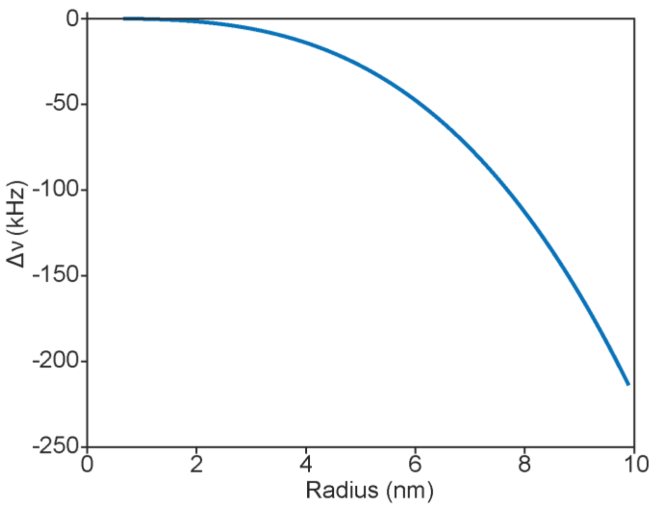

**Figure S 13.** Analytical calculation of the resonance frequency shift induced by objects of refractive index=1.43 for radii <10 nm.

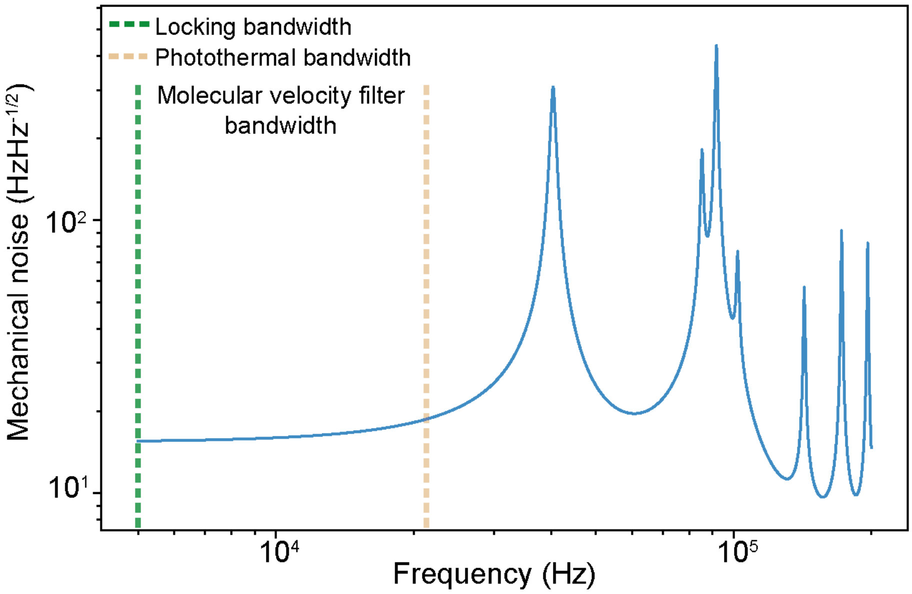

**Figure S 14.** The frequency noise spectral density for the mechanical motion of the cavity assembly extracted from finite element simulations of the mechanical modes. The resonant mechanical modes lie outside of the velocity-filter bandwidth indicating that the cavity is highly stable within the observation window. The amplitudes of the resonant mechanical modes are below the detector noise limit and are less than the calculated resonance shift for a <1 nm molecule.

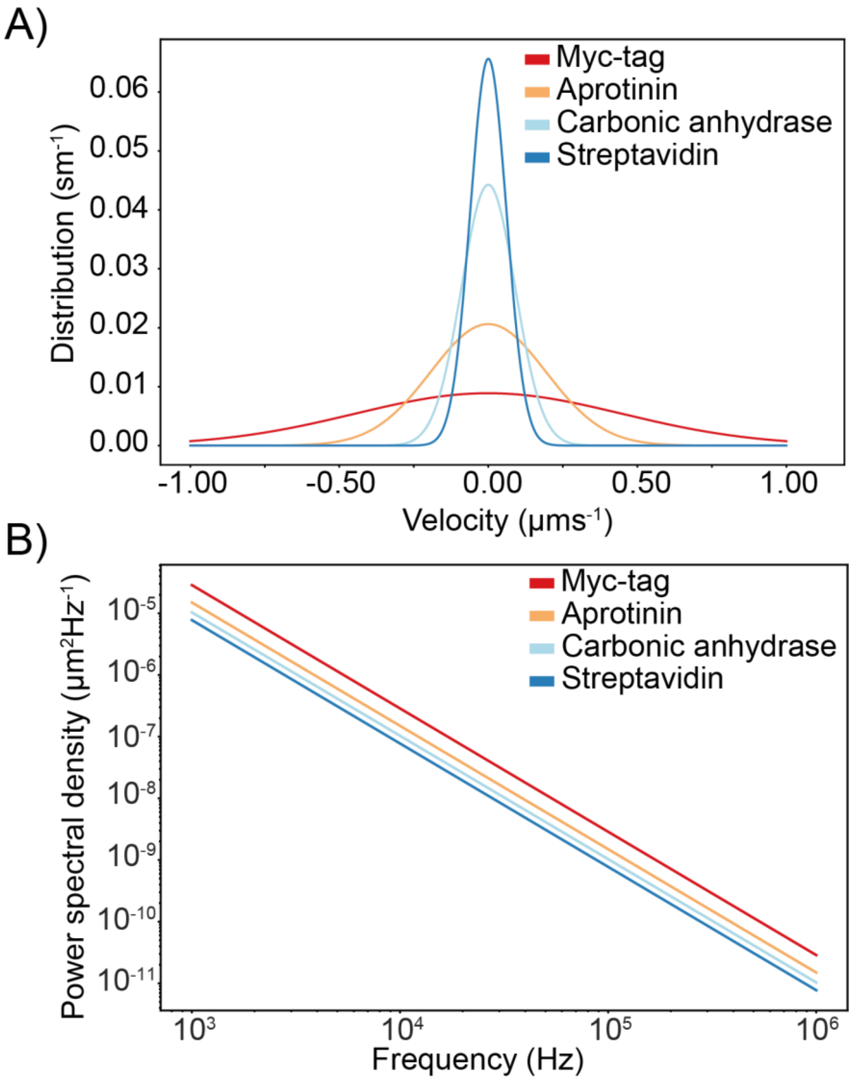

**Figure S 15 A)** Calculated velocity distribution profiles for Myc-tag, aprotinin, carbonic anhydrase and streptavidin **B)** Mean-square-displacement power spectral density plot (MSDPSD), the upper and lower bounds are replicated in the main text, Fig 5A. Integrating within the bandwidth of our velocity filter observation window (5 kHz-150 kHz) provides an approximate MSD for the molecule.

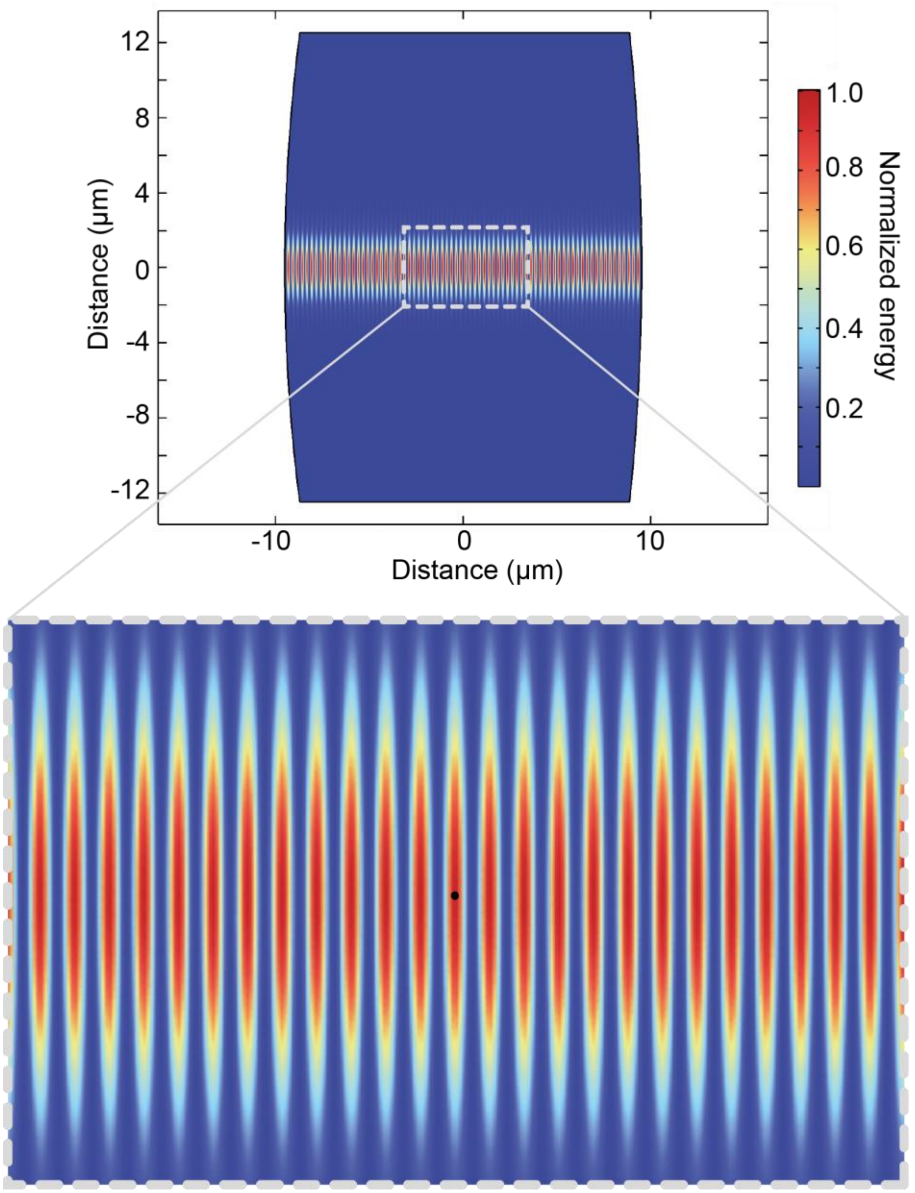

**Figure S 16.** Simulated cavity mode given the mirror properties of cavity one (Table S 4) showing the normalized square of the electric field which is proportional to the total energy stored in the cavity. The zoomed image shows a particle localized in an antinode in the center of the optical mode.

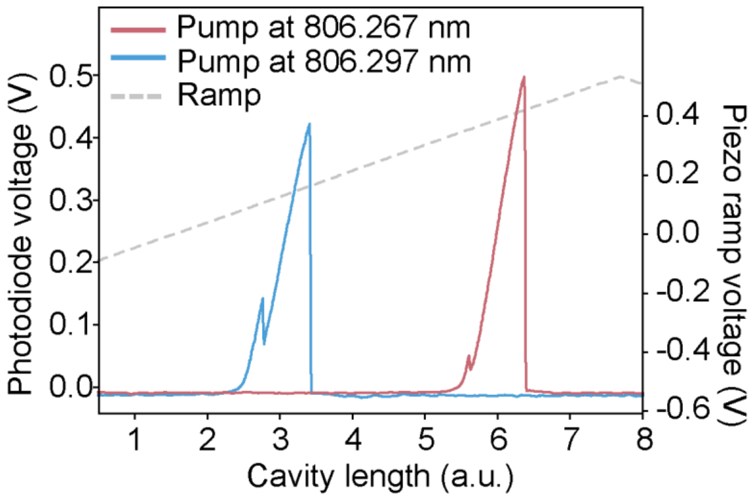

**Figure S 17.** Photothermal induced broadening of the transmitted cavity resonance in water, collected under cavity length tuning at two different pump wavelengths. The blue shift in pump wavelength shifts the resonance position to lower piezo ramp voltage, demonstrating that increased piezo ramp voltage corresponds to increased cavity length. Furthermore, the direction of the broadening is indicative of a negative thermo-optic coefficient of the medium as is expected in water. Despite the low circulating power (5.5 mW), photothermal broadening was apparent and leveraged to enable the high sensitivity of this measurement. The smaller peaks originate from polarization splitting due to the birefringence of the cavity mode. Data collected with cavity four.

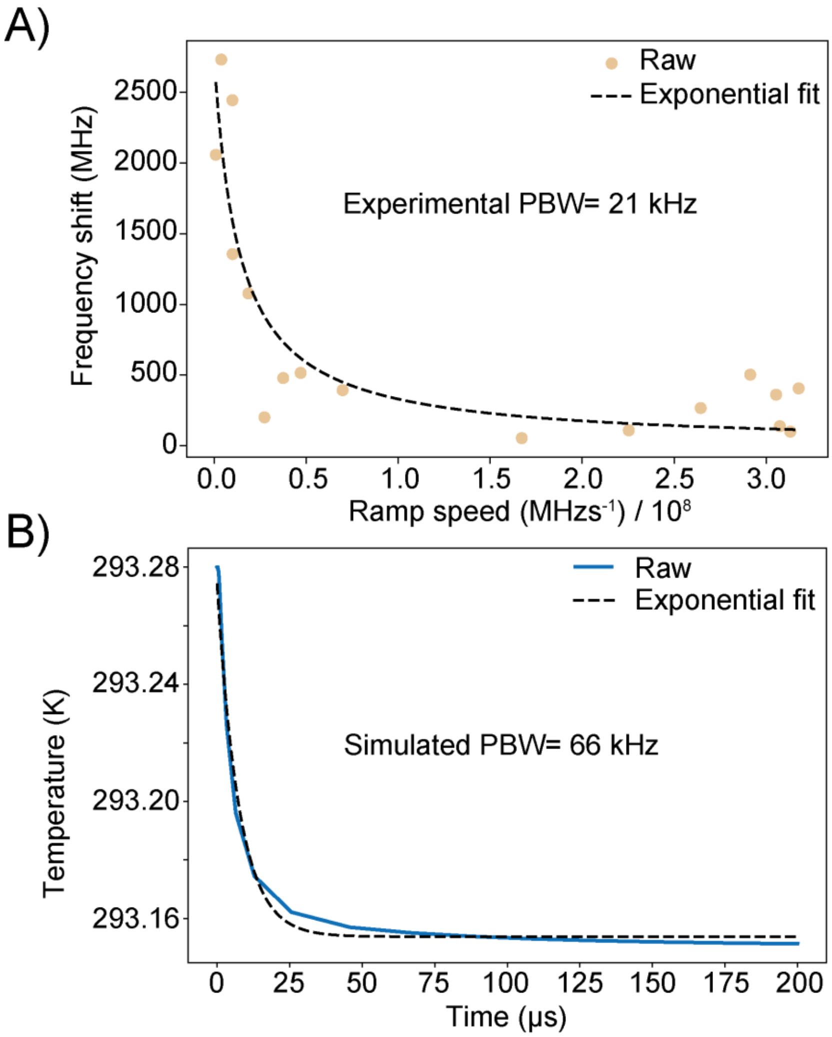

**Figure S 18. A)** Experimental determination of the photothermal bandwidth (PBW). The calculated bandwidth is 21 kHz, defining the upper limit of the molecular velocity filter (data collected with cavity four). As a comparison, **B)** finite element simulations of the rate of cooling in the cavity were performed to theoretically quantify the PBW

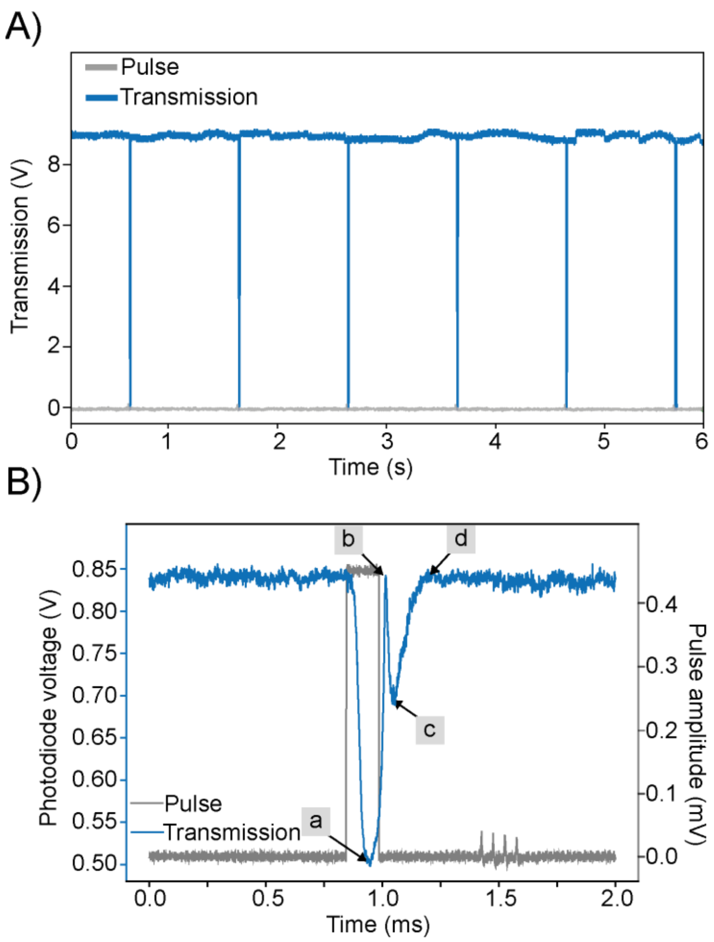

**Figure S 19. A)** Transmitted intensity of locked cavity showing perturbations to the lock when 1 mV voltage pulses are applied to the piezos at a frequency of 1 Hz. **B)** The applied pulse, input-power and cavity locking parameters can be optimized to mimic signals induced by diffusing molecules. The step down voltage (gray) produced a steep reduction of the locked transmission signal due to the photo-thermal effect (a), this was followed by a brief recovery to the locked state by the PI feedback loop (b) followed by a second descent of the transmission signal as the step-up voltage of the pulse shifts the cavity in the opposite direction (c), finally the PI control recovers the locked state (d). Data collected with cavity four.

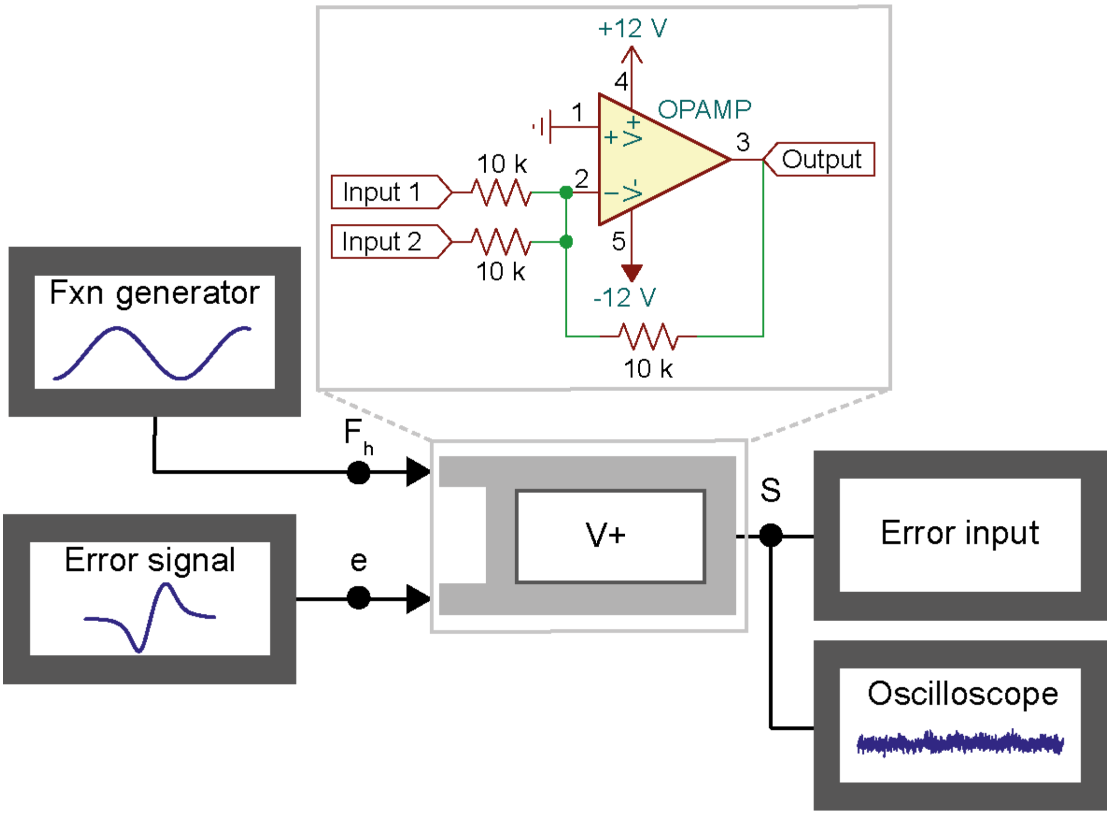

**Figure S 20.** Schematic off the apparatus used to measure the locking bandwidth of the cavity, which was measured by adding a harmonic perturbation ($F_h$) from the function generator (Fxn generator) of known frequency and amplitude together with the error signal (e) using a voltage adder (V+) to the PI input (error input).

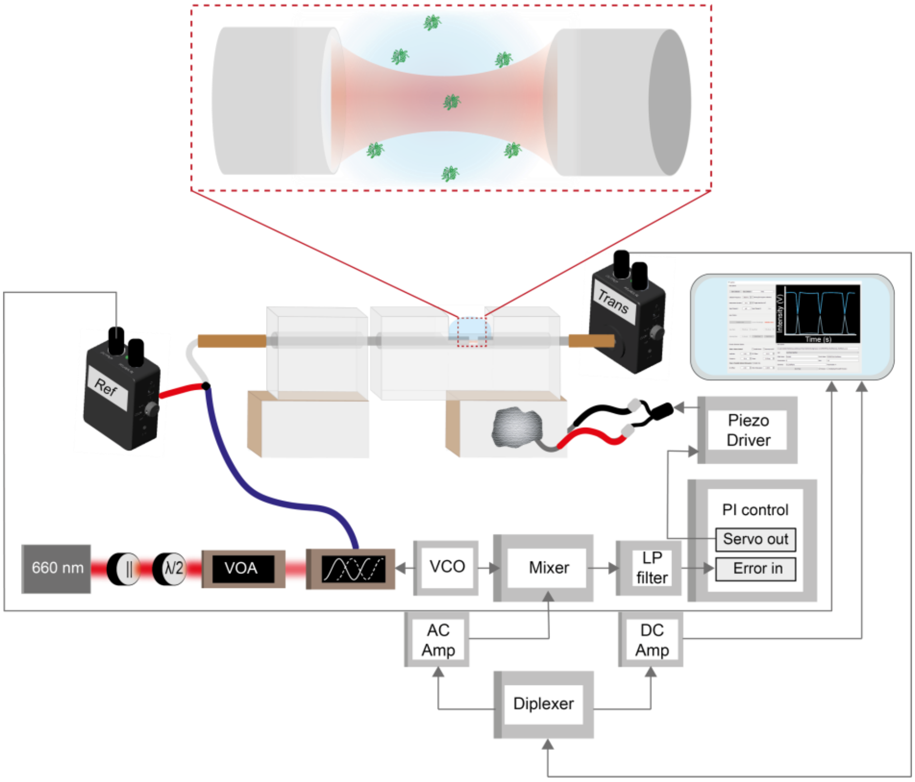

**Figure S 21.** Complete diagram showing the optical and electronic components of the setup, described in detail in the Optical setup section of the Materials and Methods**.**

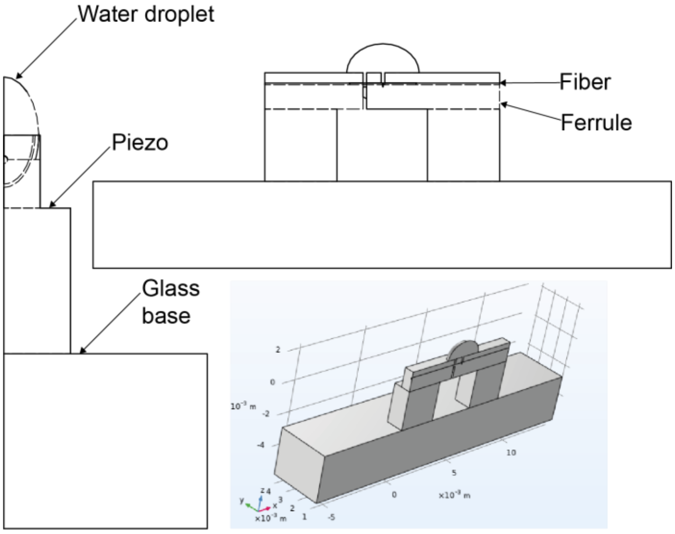

**Figure S 22.** Schematic of the cavity assembly model used for fiber mechanics COMSOL simulations**.**
